# Stress fibers orient traction forces on micropatterns: A hybrid cellular Potts model study

**DOI:** 10.1101/2022.04.18.488715

**Authors:** Koen Schakenraad, Gaia I. Martorana, Bente H. Bakker, Luca Giomi, Roeland M.H. Merks

## Abstract

Adhering cells exert traction forces on the underlying substrate. We numerically investigate the intimate relation between traction forces, the structure of the actin cytoskeleton, and the shape of cells adhering to adhesive micropatterned substrates. By combining the Cellular Potts Model with a model of cytoskeletal contractility, we reproduce prominent anisotropic features in previously published experimental data on fibroblasts, endothelial cells, and epithelial cells on adhesive micropatterned substrates. Our work highlights the role of cytoskeletal anisotropy in the generation of cellular traction forces, and provides a computational strategy for investigating stress fiber anisotropy in dynamical and multicellular settings.

**Author summary:** Cells that make up multicellular life perform a variety of mechanical tasks such as pulling on surrounding tissue to close a wound. The mechanisms by which cells perform these tasks are, however, incompletely understood. In order to better understand how they generate forces on their environment, cells are often studied *in vitro* on compliant substrates, which deform under the so called “traction forces” exerted by the cells. Mathematical models complement these experimental approaches because they help to interpret the experimental data, but most models for traction forces on adhesive substrates assume that cells contract isotropically, i.e., they do not contract in a specific direction. However, many cell types contain organized structures of stress fibers - strong contracting cables inside the cell - that enable cells to exert forces on their environment in specific directions only. Here we present a computational model that predicts both the orientations of these stress fibers as well as the forces that cells exert on the substrates. Our model reproduces both the orientations and magnitudes of previously reported experimental traction forces, and could serve as a starting point for exploring mechanical interactions in multicellular settings.

## 1 Introduction

The mechanical properties of the environment play a crucial role in many cellular processes, such as stem cell differentiation [1, 2], durotaxis [3, 4] and protein expression [5]. Conversely, cells mechanically influence their environment by applying traction forces, with direct biomedical consequences, as in cancer metastasis [6–8] or asthma [9]. These traction forces are intimately related with the cell shape [10–14] and the presence of actin stress fibers [13, 15, 16]. For instance, the total traction [11, 17–20] and the total amount of mechanical work [3, 14, 21, 22] that the cell applies on the substrate increase with the cell spreading area.

The distribution of traction forces within the cell has also been extensively studied, showing, e.g., that traction forces are larger further away from the centroid of the cell [23, 24] and accumulate at the cell periphery [25, 26], as a consequence of contractility throughout the whole cell [27]. Many studies focus on the magnitudes of the traction forces only [8, 20, 28–30], but the directions of these forces are important for understanding, for instance, cell migration [31]. Many theoretical models consider cell contractility to be isotropic and homogeneous [14, 20, 30, 32, 33], but cannot describe the anisotropic contractility caused by actin stress fibers, which strongly influence the direction of traction forces [13, 15, 16]. Because of this, in many contractility models traction forces are always normal to the surface [12, 14, 34–36], although experimental studies on micropatterns [14, 15] and during cell spreading [18] clearly show that traction forces have significant tangential components. Other models take the opposite limit in which the local cell shape and stress fiber orientation are not taken into account and traction forces generally point to the centroid of the cell [24, 32].

The work in this paper overcomes these limitations using a hybrid model that combines a finite difference method for the cytoskeleton based on liquid crystal (LC) theory, which some of us developed in Ref. [16, 37], with a Cellular Potts Model (CPM) for cell shape. Using this hybrid LC-CPM method, we calculate the distribution and orientation of traction forces at the cell periphery by taking into account both the local cell shape and the structure of the actin cytoskeleton. We theoretically study single cells adhering to adhesive micropatterns [38], and find qualitative agreement between our simulations and previously published experimental data on fibroblasts, endothelial cells, and epithelial cells, demonstrating the importance of cytoskeletal anisotropy in determining the magnitudes and orientations of cellular traction forces.

To understand the complex interplay between cellular mechanics and geometry, cells are often studied *in vitro* on an adhesive substrate [39]. Traction forces can be measured using microfabricated elastomeric pillar arrays [17, 40, 41] or traction force microscopy [42–44]. The latter is often combined with micropatterned substrates [38] to ensure reproducible cell shapes which allows for a more systematic analysis of the data. On stiff substrates, most animal cells spread out and adhere to the substrate via focal adhesions [45] mainly lying along the cell contour. This results in flat cells with a large area that exert mainly contractile forces [39]. Several types of mathematical models have been developed to interpret such experimental findings and predict the shapes of adherent cells [39]. The simplest among these are two-dimensional contour models [10, 16, 46–50], in which the cell is completely described by a closed curve that represents the cell contour. For adherent cells with a limited number of discrete adhesion sites, for instance cells on micropillar arrays, the contour consists of a set of connected arcs, hereafter referred to as “cellular arcs”, which connect two consecutive adhesion sites. Contour models predict the shapes of these cellular arcs as a function of the forces in the cytoskeleton and along the contour.

The simplest contour model, often referred to as the “Simple Tension Model” (STM), describes the total force per unit length along the cell contour, ***f***_tot_, originating from the tension in the actin cortex, *λ****T***, and the isotropic contractile stress of magnitude, in the two-dimensional situation given by an effective surface tension, *σ*, in the bulk of the cell [10, 46, 47],

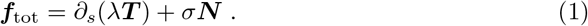

Here, ***T*** is the tangent unit vector along the cell contour, ***N*** the inward pointing normal unit vector, and *∂*_*s*_ is a derivative with respect to the arc length *s*. Here, *σ >* 0 models the isotropic contractility of the internal cytoskeleton, and *λ* describes the contractile forces arising from myosin activity in the cell cortex. The term *∂*_*s*_(*λ****T***) describes the net force per unit length on the cell contour originating from spatial variations in the cortical tension *λ* or in the orientation ***T*** of the cell contour. Analogously to the Young-Laplace equation of capillarity, Eq. 1 implies that at equilibrium, *i*.*e*., when ***f***_tot_ = **0**, the balance between bulk contractility and line tension leads to concave circular arcs with radius *R* = *λ/σ* [10, 46, 47]. *Extensions to* the STM include an elastic contribution to the line tension [10, 47], bending elasticity of the cell membrane [33, 48], and anisotropic bulk contractility [16, 37], which we discuss below.

Previous work [16, 37] extended the existing contour models by incorporating a directed, anisotropic, contractile bulk stress into the Simple Tension Model. This anisotropic contractility originates from actin stress fibers [51, 52], which we treat as contractile force dipoles with average local orientation θ_SF_ [53, 54]. Together with the isotropic contribution [Eq (1)] this leads to an overall force per unit length along the contour given by

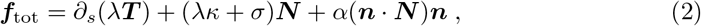

with ***n*** = (cos *θ*_SF_, sin *θ*_SF_) a unit vector parallel to the average stress fiber orientation, *κ* the local curvature of the cell boundary, *α >* 0 the magnitude of the directed contractile stresses, and where we used *∂*_*s*_***T*** = *κ****N***. Here, *α* is treated as a constant, but in Section 2.2 we will use the language of nematic liquid crystals to make *α* proportional to the local degree of alignment between the stress fibers. Upon again solving ***f***_tot_ = **0**, the equilibrium shape of the concave cellular arcs is found, in the presence of anisotropic contractility, to be given by a segment of an ellipse with aspect ratio 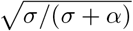 and whose longitudinal direction is parallel to the local orientation of the stress fibers *θ*_SF_. These predictions were experimentally verified by an analysis of the shapes of anisotropic fibroblastoid (GDβ1, GDβ3) and epithelioid (GEβ1, GEβ3) cells [16]. Because of the discrete nature of the adhesion sites in this contour model, cells are always concave and comparisons to experimental data are limited to cells adhering to substrates at a small number of adhesion sites.

In recent years, the Cellular Potts Model (CPM), best known for studies of multicellular systems (see, e.g., Refs. [55–57]), has emerged as a powerful computational framework for studying the shape [12, 58, 59] and motility [60–62] of single cells. In the CPM, the cell is represented by a collection of occupied lattice sites on an often two-dimensional square lattice, where a ‘lattice site’ here is used to indicate the “pixels” in a regular, square grid. The dynamics of the CPM is governed by a Monte Carlo method via the Metropolis algorithm [63]. During each step of the simulation, the state (occupied or not occupied) of a randomly chosen lattice site is copied into a random neighbor lattice site. Using a predefined energy functional, the Hamiltonian *H*, the energy change Δ*H* as a result of the copy is calculated. Then, the copy is accepted if Δ*H <* 0 and accepted with probability *e*^*−*Δ*H/µ*^ if Δ*H >* 0, where the reference energy *µ* is called the “motility energy” (aka “cellular temperature”, assuming a suitable prefactor) and is a measure for the random activity of the cell. Each term in the Hamiltonian describes a particular cellular force field ***F*** via the relation ***F*** = *−∇H*, which can be calculated for every configuration of the cell [36].

Albert and Schwarz [12] used a CPM approach to apply the Simple Tension Model, as well as the more extended “tension-elasticity model” [10, 47], to cells adhering to micropatterns of arbitrary shape. Here, we focus on the STM, for which the Hamiltonian is given by

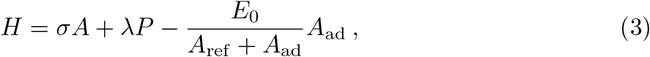

with *A* the cell area, *P* the length of cell perimeter, and *A*_ad_ represents the area of the adhesive pattern covered by the cell. Minimizing of the first two terms in Eq (3) is equivalent to letting the system evolve under the force field given in Eq (1) [10]. The last term represents the adhesion energy of the cell with the substrate, with strength *E*_0_. Because the number of adhesion molecules in a cell is finite, the adhesion energy saturates with the covered area as determined by the reference area *A*_ref_.

Here, we develop an anisotropic hybrid liquid crystal-Cellular Potts Model (LC-CPM) framework to apply the contour model in Eq (2) to cells adhering to micropatterns. The paper is organized as follows: in Section 2.1 we theoretically study how stress fibers affect cell shape, in Section 2.2 how cell shape affects the orientations of stress fibers, and in 2.3 we study the interplay between shape and cytoskeleton. We demonstrate that our results are consistent with earlier analytical and numerical approaches, and our simulations reproduce cell shape and orientations of stress fibers of previously published experimental data on fibroblasts, endothelial cells, and epithelial cells on adhesive micropatterned substrates. In Section 2.4, we predict how the anisotropy of stress fibers affects the traction forces that the cell boundary exerts on the adhesive substrate, and we reproduce prominent anisotropic features in experimentally observed traction force patterns of fibroblasts, endothelial cells and epithelial cells.

## 2 Results

### 2.1 Cytoskeletal organization controls cell shape

Following Albert and Schwarz [12], we apply the anisotropic tension model to micropatterns of arbitrary shape by implementing it in the Cellular Potts Model. In order to do so, we need to translate the anisotropic contractile force density ***f***_*a*_ = *α*(***n · N***)***n*** to an energy description. However, unlike the forces in Eq (1), the force considered here is derived from inherently non-equilibrium processes [53, 54] and is a non-conservative, *active* force. Consequently, ***f***_*a*_ cannot be derived from a Hamiltonian. For a review on non-equilibrium forces and active matter, see, e.g., Ref. [64]. Instead, we follow previous work that has incorporated other non-equilibrium processes, such as chemotaxis and durotaxis, in the Cellular Potts Model by calculating directly the mechanical work associated with a CPM copy: Δ*H*_*a*_ = *− W*_*a*_ (see, e.g., [65, 66]). For a given displacement of the cell boundary, i.e., the addition or removal of one lattice site from the cell, the total work *W*_*a*_ is obtained by integrating the force density ***f***_*a*_ over a displacement **d*r*** (to obtain the work per unit length along the boundary) and over a distance d*s* along the cell boundary:

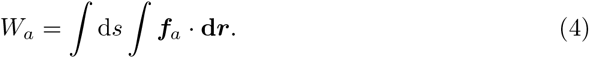

As only normal displacements of the boundary change the cell shape, we write **d*r*** =*−* d*r****N***, where d*r >* 0 if the displacement is opposite to ***N***, i.e., when the boundary moves outwards. Upon further assuming that ***n*** and ***N*** change slowly on the scale of the CPM lattice site that is added or removed, we can take them out of the integrals to find

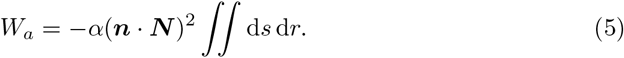

Finally, given that the displacements d*r* (along *−* ***N***) and d*s* (along ***T***) are perpendicular, the double integral is equal to the change in area of the cell, or equivalently the area covered by one lattice site, ∬ d*r*d*s* = Δ*A*, which yields

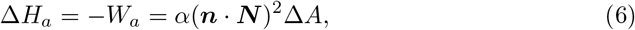

with ***n*** and ***N*** evaluated at the boundary lattice site of the cell that is copied (in case of an extension) or that is copied into (in case of a retraction), and where Δ*A* = *−* 1 lattice unit for an extension and Δ*A* = −1 lattice unit for a retraction.

To obtain the total energy change for a given CPM copy, we introduce two further simplifications. First, we assume the line tension to be constant (i.e., *∂*_*s*_*λ* = 0), although previous work has shown that *λ* can in general be different between different cellular arcs [10, 47] or even vary within a single arc [16, 37]. Second, we take the limit in which the reference area *A*_ref_ [Eq (3)] is much larger than the adhesion area, such that the adhesion energy is simplified to *γ*_ad_*A*_ad_, with *γ*_ad_ *<* 0 a negative surface tension of the adhesion area. The total energy change for a copy is then given by

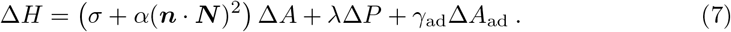

See Section 4 in the Methods section for the numerical methods used for determining the normal vector ***N*** and the length of the perimeter *P*.

Before we study cells on micropatterned substrates, we first test the validity of our CPM model by comparing the shapes of non-adhering (i.e., *γ*_ad_ = 0) single cellular arcs to previously published analytical predictions [10, 16, 46, 47]. To do so, we define as initial condition a rectangular “cell” to occupy all lattice sites in the lower half of the simulation space. This is schematically illustrated in Fig 1a, where the lattice sites that are part of the cell are displayed in blue. Then, we define two adhesion sites (green circles in Fig 1a), placed 100 lattice sites apart, where the cell is pinned to the substrate. All lattice sites that are not in between the adhesion sites (grey areas) are frozen and are not allowed to be updated during the simulation. The region in between the adhesion sites then forms a cellular arc whose shape we study.

**Fig 1.**
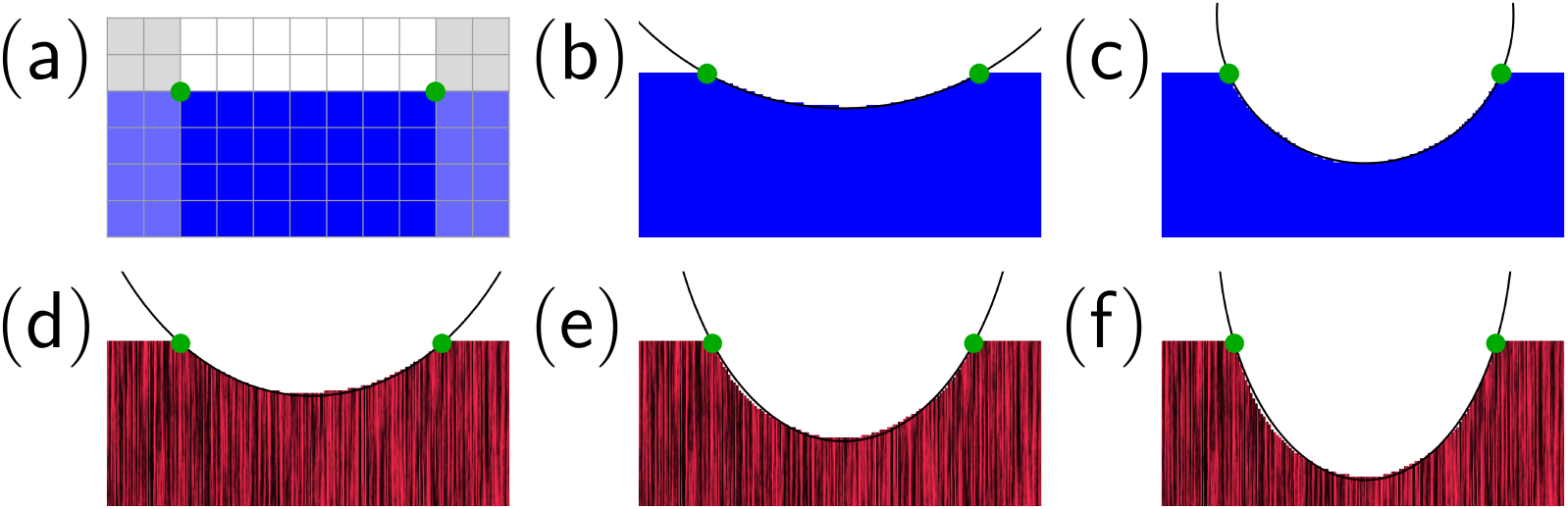
Single cellular arcs are well described by circles and ellipses. (a) Schematic of the simulation setup for studying the shape of a single arc. The initial condition is a rectangular “cell” (blue lattice sites) that occupies the lower half of the lattice sites in the simulation. Two adhesion sites (green circles), placed 100 lattice sites apart, pin the cell to the substrate. In practice, all lattice sites that are not in between the adhesion sites (grey areas) are frozen and are not updated during the simulation. A Cellular Potts Model (CPM) simulation is then performed using the Hamiltonian in Eq (7), with *γ*_ad_ = 0, to find the shape of the arc. (b,c) Configurations of single arcs for *µ/λ* = 1*/*10 lattice sites and *α* = 0 are well approximated by circles. The theoretical radii of the circles, *R* = *λ/σ*, are 50 and 100 lattice sites respectively, whereas the lattice sites. (d-f) Configurations of single arcs for *α* ≠ 0 are well approximated by ellipses. The black lines in the cell interior represent the vertically oriented stress fibers (i.e., *θ*_SF_ = *π/*2 and ***n*** = ***ŷ***). The approximating ellipses have aspect ratios given by 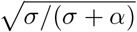 and are vertically oriented, as theoretically predicted. The lengths of the major semi-axes are given by: (d) 134 lattice sites (approximation) versus 120 lattice sites (theory), (e) 119 lattice sites (approximation) versus 120 (theory), (f) 98 lattice sites (approximation) versus 100 (theory). Parameters are given by: (d) *α* = *σ, λ/σ* = 120 lattice sites, and *µ/λ* = 1*/*12 lattice sites, (e) *α* = 2*σ, λ/σ* = 120 lattice sites, and *µ/λ* = 1*/*12 lattice sites, and (f) *α* = 2*σ, λ/σ* = 100 lattice sites, and *µ/λ* = 1*/*10 lattice sites.

First, we study a cell with an isotropic cytoskeleton, i.e., *α* = 0. The Hamiltonian then reduces to *H* = *σA* + *λP*, which is equivalent to a force per unit length along the contour given by Eq 1 [10]. Hence, we expect to find a concave circular arc with radius *R* = *λ/σ* [10, 46, 47]. In Figs 1b,c we show the results of two simulations with the same line tension λ, but with different values of the isotropic bulk stress *σ*. As expected, the arcs are concave and the arc with the larger bulk stress (Fig 1c) is more curved. The arcs are well approximated by circles (black lines in Figs 1b,c), as was also observed before for this Hamiltonian in Ref. [12]. The radii of the circles shown in Figs 1b,c are, however, slightly larger (5%-10%) than expected based on the theoretical prediction *R* = *λ/σ*. This slight underestimation of the observed radius is consistently found for all choices of the parameters *λ* and *σ*. We hypothesize that this is a consequence of random fluctuations in the CPM due to the finite motility energy *µ*. These random fluctuations favor states of the system that can be realized in many different ways (i.e., shapes with large *entropy*), leading to slightly less curved cellular arcs, and consequently bigger circles than expected based on the energy minimization.

Next, we include the directed bulk stress (i.e., *α* ≠ 0), and assume a predefined and constant orientation of the cytoskeleton ***n*** = ***ŷ*** (i.e, *θ*_SF_ = *π/*2). The resulting Hamiltonian [first two terms in Eq (7)] is equivalent to a force per unit length described by the second and third term of Eq (2). Hence, we expect the cellular arcs to approximate a segment of an ellipse with aspect ratio 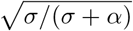 and whose long semi-axis of size *λ/σ* is oriented vertically, thus parallel to the stress fibers [16, 37], although we do not expect the agreement to be perfect because we ignore the first term in Eq (2) by assuming the line tension *λ* to be constant. Figs 1d-f show the cellular arcs together with segments of ellipses matching the arcs. The thin black lines in the interior of the cells visualize the vertical orientation of the stress fibers. Fig 1d shows a cellular arc for *σ* = *α* and *λ/σ* = 120 lattice sites. In Fig 1e the directed bulk contractility is increased to *α* = 2*σ* resulting in a more curved shape and a narrower (larger aspect ratio) ellipse. In Fig 1f the line tension is additionally decreased such that *λ/σ* = 100 lattice sites, resulting in an even more curved shape and a smaller ellipse than in Fig 1e. The orientations and aspect ratios of the ellipses shown in Figs 1d-f match the exact solution, but, similar to what we observed for the circles, the ellipse predicted by the numerical solution in Fig 1d is slightly larger than the exact solution. We stress that the accuracy of the numerical solution increases with the ratio *α/σ*. This behavior originates from the fact that directed stresses, embodied in the parameter *α*, increase the overall contractility of the cell, thereby requiring a higher cortical tension to prevent the cell from collapsing. Since *λ* is treated as a constant in the present CPM this feedback mechanism cannot take place, resulting into higher bending of the cellular arcs and thus smaller ellipses. This effect compensates for the fact that both ellipses and circles are larger than expected because of the inherent randomness of the CPM.

Summarizing, although the correspondence between the theoretical predictions [10, 16, 37, 46, 47] and our simulations on the shapes of single cellular arcs is not exact, the lattice-based CPM approximates the continuous curves predicted by the theory remarkably well. For a numerical treatment of these models that explicitly describes the cell boundary, and therefore produces even more accurate shapes, we refer to Ref. [37]. Here, we take advantage of the possibilities of the CPM to study cells adhering to continuous micropatterned geometries in Sections 2.2, 2.3, and 2.4.

### 2.2 Cell shape controls cytoskeletal organization

In order to apply the method introduced in Section 2.1 to entire cells, we require a theoretical description of the cytoskeleton to define the non-constant stress fiber orientation ***n*** at every lattice site. For this purpose we employ a continuous phenomenological model of the actin cytoskeleton, using the language of nematic liquid crystals [67] developed in Refs. [16, 37]. We emphasize, however, that our method of anisotropic cytoskeletal contractility in the CPM as summarized by Eq (6) can in principle be combined with any theoretical description of the cytoskeleton (e.g., Ref. [68]).

As detailed in Ref. [37], the configuration of the stress fibers in the actin cytoskeleton is described by the two-dimensional nematic tensor, whose Cartesian representation is given by:

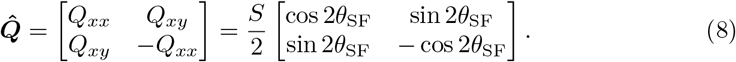

where 0 ≤ *S* ≤ 1 is the so called nematic order parameter. *S* measures the amount of orientational order of the stress fibers, where *S* = 1 represents perfect local alignment between stress fibers, and *S* = 0 represents randomly oriented stress fibers. This model does not account for the spatial variations in the actin density observed within the cell [14, 69–71].

The general idea behind our cytoskeleton model is based on the experimental observation [15, 16, 72–75] that stress fibers in highly anisotropic cells preferentially align with each other and with the cell’s edges. For an overview of the physical, chemical, and biological mechanisms that are possibly involved in these aligning interactions, see Ref. [37]. The trend is phenomenologically captured by the Landau-de Gennes free-energy *F*_cyto_ [67]:

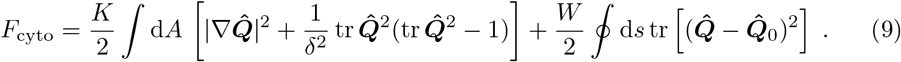

The first integral in Eq (9) is responsible for the alignment of stress fibers with one another, as it penalizes gradients in the orientation of the stress fibers and in their agree of alignment (first term). The constant *K* expresses the stiffness of the average stress fiber orientation ***n*** with respect to splay and bending deformations. The second term equals *S*^2^(*S*^2^*/*2*−* 1)*/*(2*δ*^2^), which favors perfect orientational order (i.e., *S* = 1), where the length scale *δ* sets the size of associated boundary layers in which the order parameter *S* deviates from its preferential value. In particular, *δ* determines the typical size of regions where the order parameter *S* drops to zero to compensate for a strong local gradient in the stress fiber orientation *θ*_SF_. The second integral is the

Nobili-Durand anchoring energy [76], expressing the energetic cost of associated with a departure of 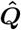 from the peripheral configuration

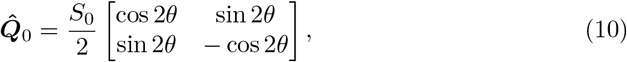

where *θ* is an angle parallel to the cell edge, such that ***T*** = (cos *θ*, sin *θ*). In addition to stress fibers aligning to the cell edge, we assume that they do so with large nematic order: *S*_0_ = 1. The phenomenological constant *W >* 0 [Eq (9)] measures the strength of the parallel anchoring, i.e., it measures to what extent stress fibers align with the cell edge. For a description of the numerical methods employed for minimizing the energy, see Section 4.

As we mentioned in Section 1, the directed bulk contractility *α* [Eq (2)] increases with the local alignment between stress fibers. Having introduced the nematic order parameter *S*, we now make this relation quantitative by setting *α* = *α*_0_*S*, with *α*_0_ a constant (see also Ref. [37]). Apart from this, we treat the contractility parameters *α*_0_, *σ*, and *λ*, which model the tension at the cell edge, independently from the parameters *K, W*, and *δ*, which model the interior of the cell.

We combine this model for the cytoskeleton with the Cellular Potts Model for cell shape described in Section 2.1. We alternatingly update the CPM, using the Hamiltonian given by Eq (7) with *γ*_ad_ ≠ 0, and the structure of the cytoskeleton by minimization of the free energy *F*_cyto_ [Eq (9)], until both the cell shape and the cytoskeleton reach a steady state. For details, see Section 4. Before we study the interplay between cytoskeleton and cell shape in Section 2.3, in this section we illustrate how cell shape affects the cytoskeleton and compare our model predictions to previously published experimental data. Fig 2 shows the results of simulations performed on three convex micropatterns (Fig 2a), namely a rectangle of aspect ratio 2, a stadium-shape of aspect ratio 2 and a circle. For cells adhering to these convex patterns, all contractile forces described by the first two terms in the Hamiltonian of Eq (7) are directed inwards. Hence the steady-state shape of these cells is, independent of the exact values of *α*_0_, *σ*, and *λ*, identical to the convex micropatterns that they adhere to. This makes these shapes well suited for studying the effect of cell shape on cytoskeletal organization. In Fig 2b we study the cytoskeleton for increasing parallel anchoring of the stress fibers with the cell edge. The importance of this parallel boundary anchoring, quantified by *W*, relative to parallel alignment in the bulk, quantified by *K*, is described by the *anchoring number An*, a dimensionless number which we introduced in Ref. [37] and is given by

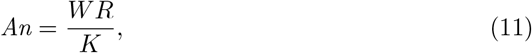

with *R* a typical length scale in which the cytoskeleton is confined, see Fig 2a. For *An «* 1 bulk elasticity dominates boundary anchoring, resulting in a uniformly oriented cytoskeleton with large deviations form parallel anchoring with the cell’s edges. On the other hand, for *An »* 1, anchoring dominates, leading to perfect alignment of the stress fibers with the boundary.

**Fig 2.**
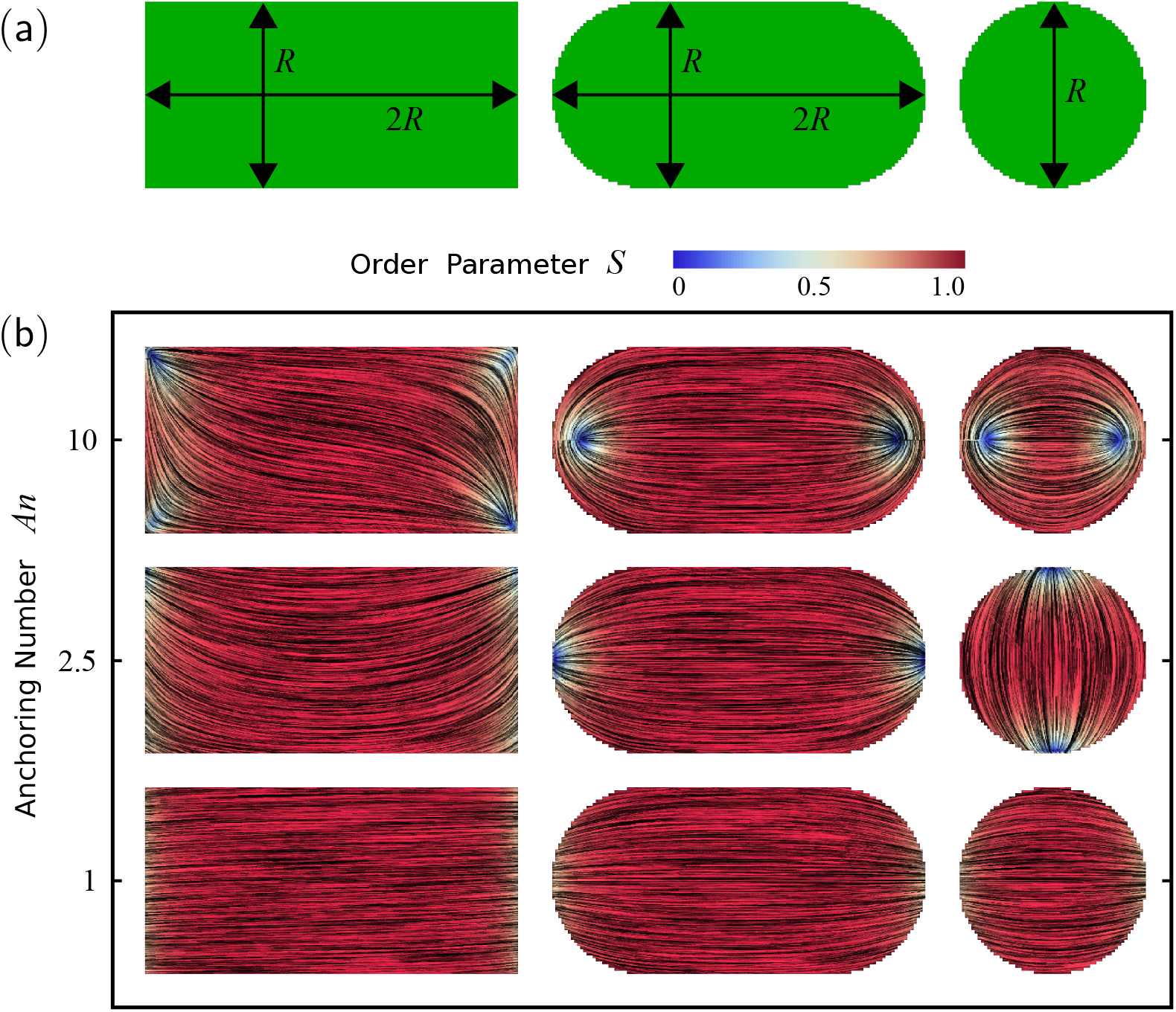
Cytoskeletal structure of cells on adhesive micropatterns with convex shapes. (a) Adhesive micropatterns with the shape of a rectangle of aspect ratio 2, a stadium of aspect ratio 2, and a circle. The length scale *R* = 60 lattice sites is used to calculate the anchoring number *An*, defined in Eq (11). (b) Cells with increasing anchoring number (*An* = 1, *An* = 2.5, and *An* = 10) on the adhesive micropatterns shown in (a). The black lines in the interior of the cells represent the orientation field ***n*** = (cos *θ*_SF_, sin *θ*_SF_) of the stress fibers and the background color indicates the nematic order parameter *S*. The spatial averages of the order parameter are given, from top to bottom, by: 0.90; 0.94; 0.97 (rectangles), 0.92; 0.94; 0.97 (stadiums), and 0.81; 0.87; 0.94 (circles). For all cells, *δ* = 9 lattice sites, *λ/σ* = 300 lattice sites, *α*_0_*/σ* = 2, *µ/λ* = 1*/*30 lattice sites, *µ/*|*γ*_ad_| = 1*/*20 of the area of a lattice site, and *K*Δ*t/*(*ξ*_*r*_*R*^2^) = 2.5 *·* 10^*−*6^. For definitions of Δ*t* and *ξ*_*r*_, see Section 4.

The left column of Fig 2b shows the cytoskeleton of a cell, adhering to the rectangular micropattern, for *An* = 10, *An* = 2.5 and *An* = 1 respectively, where *R* is the short side of the rectangle (see Fig 2a). The black lines in the interior of the cells represent the orientation field ***n*** = (cos *θ*_SF_, sin *θ*_SF_) of the stress fibers and the background color indicates the nematic order parameter *S*. Consistent with our previous results on rectangular geometries [37], stress fibers align with all edges for large boundary anchoring (*An* = 10), but they align only with the rectangle’s longitudinal direction for small boundary anchoring (*An* = 1), consistently with various experimental observations on cells adhering to elongated adhesive micropatterns and adhesive stripes [14, 15, 71, 74, 75]. We note that for configurations with sufficiently large anchoring numbers (*An* = 2.5 and *An* = 10), the random initial configuration of the simulation determines, independently of the anchoring number, whether the stress fibers near the short edges of the rectangle bend in opposite directions, leading to an “S”-shaped cytoskeleton (*An* = 10 in Fig 2), or in the same direction, leading to a “U”-shaped cytoskeleton (*An* = 2.5 in Fig 2).

Next, we focus on shapes distinct from those of cells adhering to a small number of discrete adhesion sites, which we studied in Ref. [37], and investigate the actin cytoskeleton on convex micropatterns with curved edges. The middle column of Fig 2b shows the cytoskeleton, for identical values of *An* as in the left column, for a cell adhering to a stadium-shaped micropattern of aspect ratio 2, where *R* is again the length of the minor semi-axis. For *An* = 1 the cytoskeleton aligns longitudinally and the order parameter is roughly uniform and close to its perferential value *S*_0_ = 1, similarly to what we observed for the rectangle. However, unlike for rectangular patterns, the more gentle curvature of the stadium causes the stress fiber orientation at the circular caps to deflect toward the middle. This effect becomes more pronounced for *An* = 2.5, where two topological defects start to form at the ends of the circular caps, as can be seen from the more blue-shifted color. For a review on topological defects in nematic liquid crystal systems, see, e.g., Ref. [77]. For the largest *An* value the stress fibers align with the edge throughout the complete cell, causing the topological defects to move inwards and the average order parameter to decrease to 0.92. Unlike the topological defects found in the stress fiber orientation in concavely shaped cells in Ref. [37], which have topological charge − 1*/*2, these topological defects have charge +1*/*2 due to the convex cell shape. The observed cytoskeletal structures agree qualitatively with experimentally observed stress fiber distributions in fibroblasts on micropatterns of spherocylindrical shape in Ref. [14], where structures similar to our theoretical results for *An* = 10 (for relatively wide stadium-shapes, hence large *R* and large *An*) and for *An* = 1 (for narrow stadium-shapes, small *R* and *An*) are found. In Section 2.4, where we study traction forces, we compare in more detail with the experimental data in Ref. [14].

Finally, we decrease the aspect ratio to 1 while keeping the micropattern curvature constant. The right column of Fig 2b shows the cytoskeleton for a cell adhering to a circular micropattern (with *R* the diameter of the circle). The resulting configurations are qualitatively similar to those observed for the stadiums-shapes: for *An* = 1 the stress fibers align largely uniformly with small deflections inwards (average order parameter is 0.94), for *An* = 2.5 the structure bends more and topological defects start to form near the edge (average order parameter 0.87), and for *An* = 10 the stress fibers align with the cell edge everywhere, causing the defects to move inwards (average order parameter 0.81). The most important difference between circles and stadiums is that the main orientation is no longer biased toward a specific direction due to the symmetry of the shape, although the symmetry of the square lattice of the Cellular Potts Model appears to favor either horizontal or vertical configurations. The linear stress fiber structure for

*An* = 1 closely resembles the actin cytoskeleton of fibroblasts found in Refs. [78, 79] and in simulations by Pathak *et al*. [68], but this linear pattern is not found for fibroblasts in Ref. [14] or for non-keratinocyte epitheliocytes or keratinocytes in Ref. [79], where mostly isotropic configurations that lack stress fibers are found. Hence, our cytoskeleton model qualitatively reproduces experimentally found stress fiber configurations for lower *An* values, but is not applicable to describe cells that have not formed stress fibers.

### 2.3 Cytoskeleton and shape interplay on micropatterns

In previous sections we studied how the cytoskeleton affects cell shape (Section 2.1) and how the shape affects the orientation of the cytoskeleton (Section 2.2). Before demonstrating that stress fibers play an important role in directing traction forces in Section 2.4, we first study the interplay between stress fiber orientation and cell shape. We focus on two frequently studied micropattern shapes: the letter V [12, 68, 69] and the crossbow [6, 12, 70], see Fig 3. We measure the magnitude of the bulk contractility of the cells with respect to their line tension using another dimensionless number introduced in Ref. [37], the *contractility number Co*:

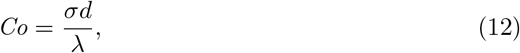

with *d* the typical distance between the ends of a non-adherent cellular arc (see Fig 3). For *Co* « 1 line tension dominates bulk contractility, resulting in straight non-adherent cellular edges, whereas *Co* » 1 leads to highly curved non-adherent cellular edges.

**Fig 3.**
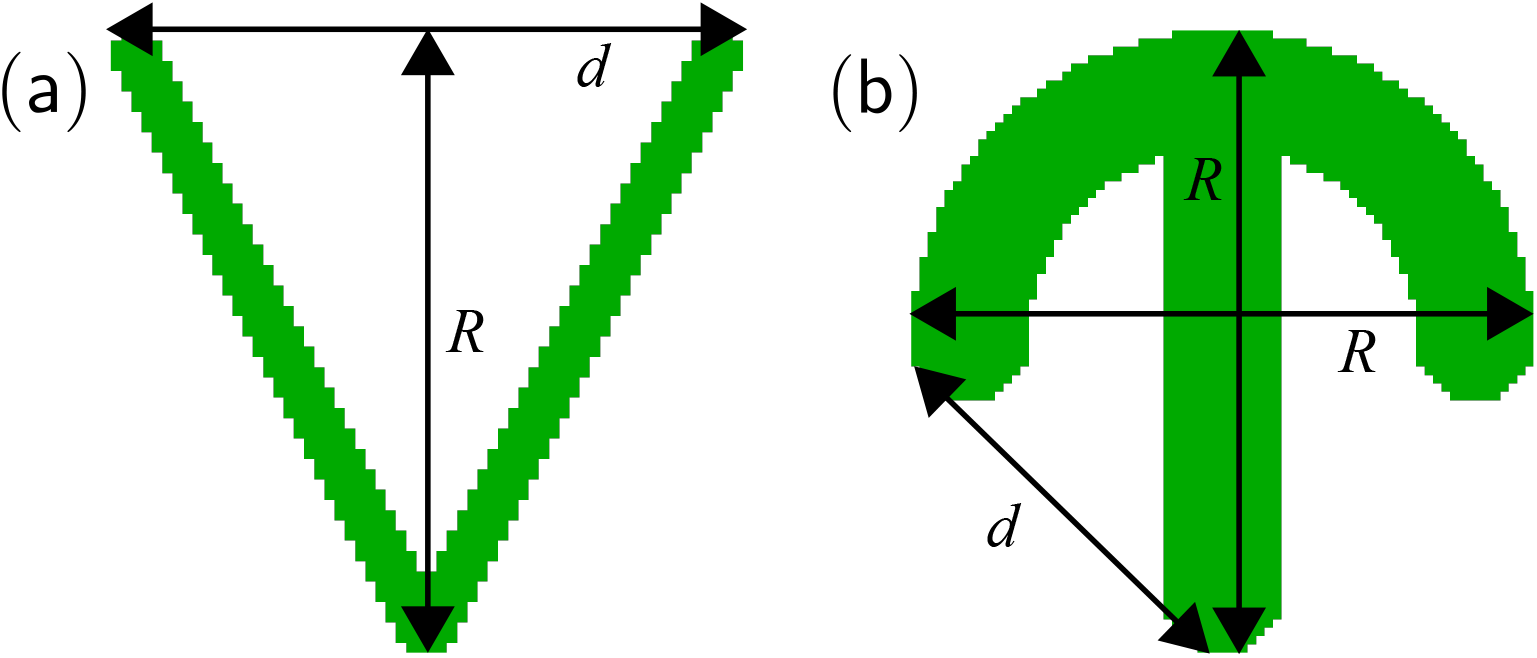
Adhesive micropatterns to study the interplay between cell shape and the orientation of the actin cytoskeleton. (a) Pattern in the shape of the letter V, where *R* = 60 lattice sites [Eq (11)] and *d* = 60 lattice sites [Eq (12)]. (b) Pattern in the shape of a crossbow, where *R* = 60 lattice sites and *d* = 42 lattice sites.

Fig 4 shows cells on the V-pattern as a function of the anchoring number *An* (on the vertical axis) and as a function of the contractility number *Co* (on the horizontal axis). For *Co* = 0, the cell has a triangular shape, whereas for nonzero *Co* the free edge at the top curves inwards. Similarly to what we observed in Fig 2b, for *An* = 10 the stress fibers align with the edge throughout the complete cell, leading to topological defects in the cell interior. Unlike for the circular and spherocylindrical patterns in Fig 2b, triangular cells feature *−*1*/*2 defects. For smaller *An*, however, the boundary anchoring is too small to bend the stress fibers sufficiently to align with all edges. Consequently, the cytoskeleton aligns in the vertical direction, leading to perpendicular alignment with the non-adherent cell edge. We note that *An* indirectly also influences the cell shape: for nonzero constant stress fiber contractility (i.e., constant *Co >* 0), increasing *An* leads to more tangential alignment of the stress fibers with the cell edge. Because the directed contractile bulk force is proportional to (***n · N***)^2^ [Eq (6)], stress fibers exert the maximal force when orthogonal to the cell edge. Consequently, increasing *An* decreases the contractile force experienced by the cell edge, as can be seen in the right column of Fig 4.

**Fig 4.**
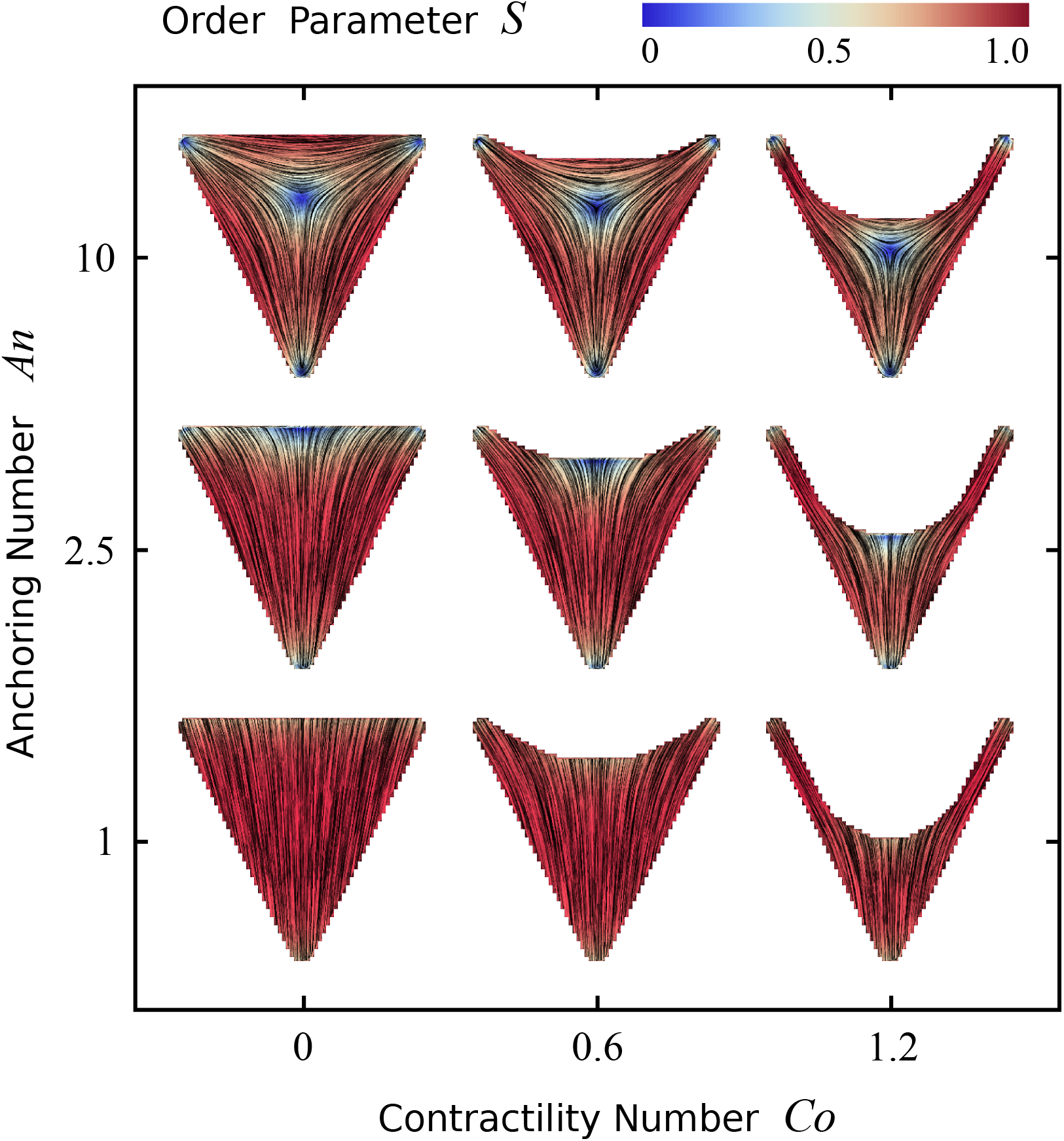
Cells on adhesive micropatterns in the shape of the letter V (see Fig 3a). The black lines in the interior of the cells represent the orientation field ***n*** = (cos *θ*_SF_, sin *θ*_SF_) of the stress fibers and the background color indicates the nematic order parameter *S*. The vertical axis corresponds to the anchoring number *An* = *WR/K* = 1, 2.5, 10 (with *R* = 60 lattice sites) and the horizontal axis to the contractility number *Co* = *σd/λ* = 0, 0.6, 1.2 (with *d* = 60 lattice sites). The spatial averages of the order parameter are given, from left to right, by: 0.76; 0.77; 0.75 (top row), 0.83; 0.85; 0.83 (middle row), and 0.93; 0.92; 0.91 (bottom row). For all cells, *δ* = 9 lattice sites, *α*_0_*/σ* = 2, *µ/λ* = 1*/*30 lattice sites, *µ/*|*γ*_ad_| = 1*/*20 of the area of a lattice site, and *K*Δ*t/*(*ξ*_*r*_*R*^2^) = 2.5 *·* 10^*−*6^. For definitions of Δ*t* and *ξ*_*r*_, see Section 4.

The result for strong anchoring and intermediate contractility best resembles the actin structures found in epithelial cells in Ref. [69] and in numerical simulations in Ref. [68]. However, an important difference is that in those studies, stress fibers align more with the non-adhesive, concave edge at the top of the pattern than with the other two edges, leading to a more horizontal actin orientation in the cell interior. This discrepancy is most likely explained by the experimental observation that stress fibers form more prominently along concave edges than along convex edges [6, 69, 71, 80], whereas our current model does not discriminate between these two scenarios. Taking this difference into account will be an important step to improve the accuracy of our cytoskeleton model in the future.

Fig 5 shows cells on the crossbow pattern (Fig 3b), again as a function of the anchoring number (vertical axis) and contractility number (horizontal axis). For small boundary anchoring, the cytoskeleton again orients vertically, leading to perpendicular alignment at the bottom and top of the crossbow shape. Conversely, for large *An*, stress fibers orient parallel to the edge throughout the cell. Interestingly, the top row of Fig 5 illustrates that *Co* indirectly influences the orientation of stress fibers: increasing the bulk contractility changes the cell shape and therefore affects the boundary conditions for the orientation of the cytoskeleton. As a result, the main orientation of the stress fibers switches from vertical to horizontal, and the two topological defects of charge +1*/*2 at opposite ends of the cell are replaced by three +1*/*2 defects surrounding a *−* 1*/*2 defect in the center. The cell shape at intermediate contractility values (middle column in Fig 5) resembles the shape of epithelial cells on a crossbow pattern found experimentally by Tseng *et al*. [6] and computationally by Albert and Schwarz [12], and the actin orientation reported in Tseng *et al*. is best reproduced by low to intermediate boundary anchoring.

**Fig 5.**
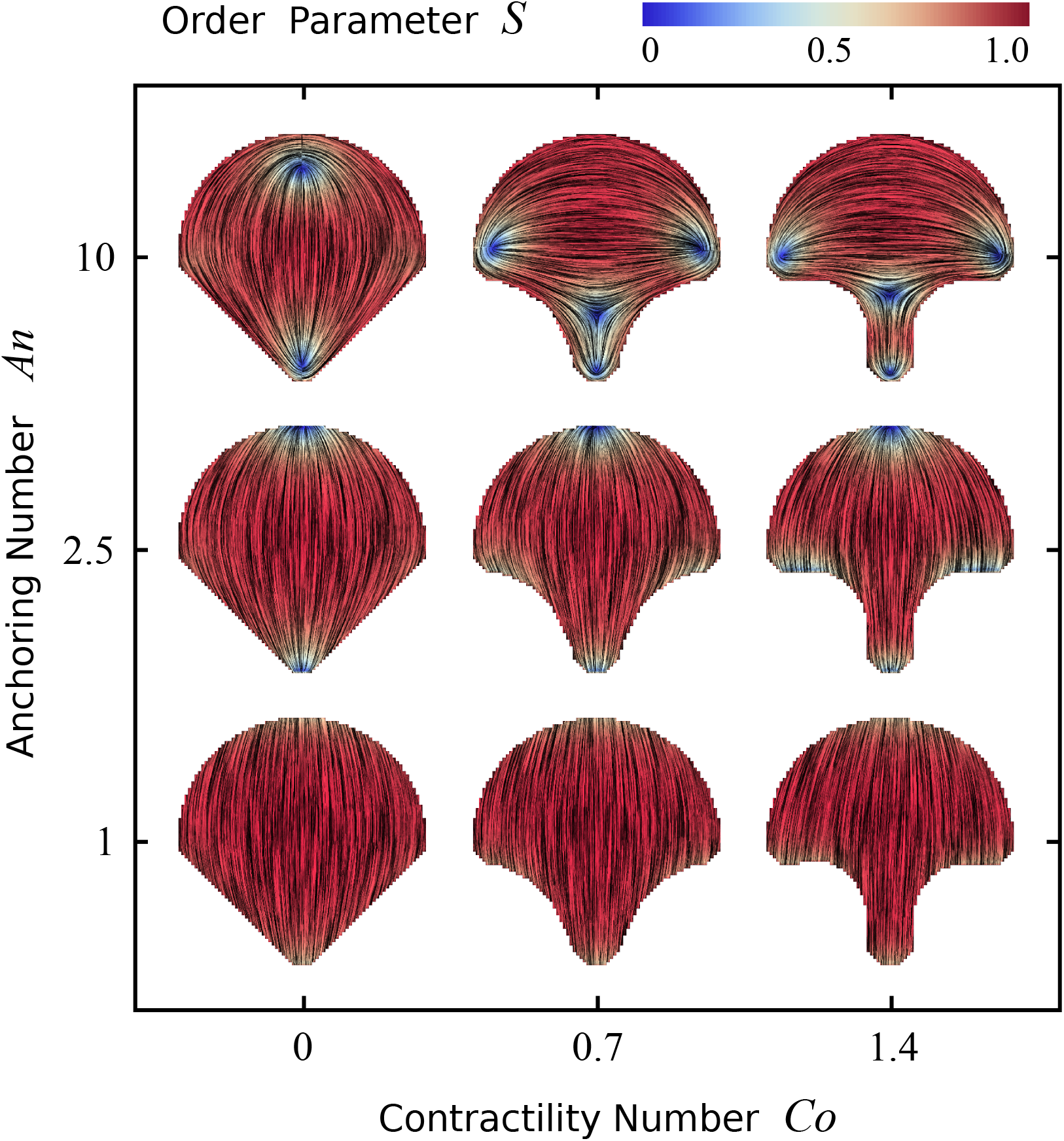
Cells on adhesive micropatterns in the shape of a crossbow (see Fig 3b). The black lines in the interior of the cells represent the orientation field ***n*** = (cos *θ*_SF_, sin *θ*_SF_) of the stress fibers and the background color indicates the nematic order parameter *S*. The vertical axis corresponds to the anchoring number *An* = *WR/K* = 1, 2.5, 10 (with *R* = 60 lattice sites) and the horizontal axis to the contractility number *Co* = *σd/λ* = 0, 0.7, 1.4 (with *d* = 42 lattice sites). The spatial averages of the order parameter are given, from left to right, by: 0.84; 0.79; 0.80 (top row), 0.89; 0.87; 0.85 (middle row), and 0.95; 0.94; 0.93 (bottom row). For all cells, *δ* = 9 lattice sites, *α*_0_*/σ* = 2, *µ/λ* = 1*/*30 lattice sites, *µ/*|*γ*_ad_| = 1*/*20 of the area of a lattice site, and *K*Δ*t/*(*ξ*_*r*_*R*^2^) = 2.5 *·* 10^*−*6^. For definitions of Δ*t* and *ξ*_*r*_, see Section 4.

### 2.4 Traction forces on micropatterns

The steady-state configurations shown in Sections 2.1, 2.2, and 2.3, are obtained when Δ*H* = 0 and the system is in mechanical equilibrium. Above micropatterned areas, the adhesive force applied by the substrate [last term in Eq (7)] balances the contractile forces generated by the cell [first two terms in Eq (7)]. The traction force per unit length that the cell edge exerts on the substrate is equal and opposite to the adhesive force per unit length to the substrate, and is given by:

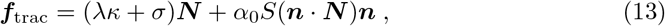

which is different from Eq (2) because here we assume the line tension *λ* to be constant (see Section 2.1). The isotropic contractility *σ* and the line tension *λ* give rise to forces normal to the cell edge [first term in Eq (13)]. However, the anisotropic contractility caused by actin stress fibers, given by the last term of Eq (13), rotates the traction forces in the direction of the local stress fiber orientation ***n***, consistent with experimental observations in Refs. [13, 15, 16]. Consequently, the traction forces in Eq (13) are not normally directed in contrast to many previous models for cell contractility [14, 34, 35]. Hence, in our model, the degree to which the traction forces deviate from the normal direction depends on the local orientation of stress fibers ***n***, the ratio between the directed contractility *α*_0_*S* and the parameters *σ* and *λ*, and on the local curvature *κ* of the cell edge. For a description of how the curvature is numerically calculated, see Section 4.

Fig 6 shows the traction forces at the cell periphery of cells on an adhesive micropattern in the shape of a stadium (see Fig 2), as a function of the anchoring number *An* (vertical axis) and the directed bulk contractility *α*_0_ (horizontal axis). Here, the micropattern shape controls the curvature *κ, An* controls the orientation of the stress fibers ***n*** (Section 2.2), and *α*_0_ sets the relative importance of directed stresses with respect to the other stresses in the cell. For *α*_0_ = 0 (left column of Fig 6), all traction forces are independent of the stress fiber orientation ***n*** and normal to the surface. At the circular caps, the larger curvature increases the effect of the line tension [Eq (13)], which increases the traction forces, reproducing findings on the relation between local curvature and traction from earlier models [12, 14, 32] and from experimental observations [11, 13, 14, 81].

**Fig 6.**
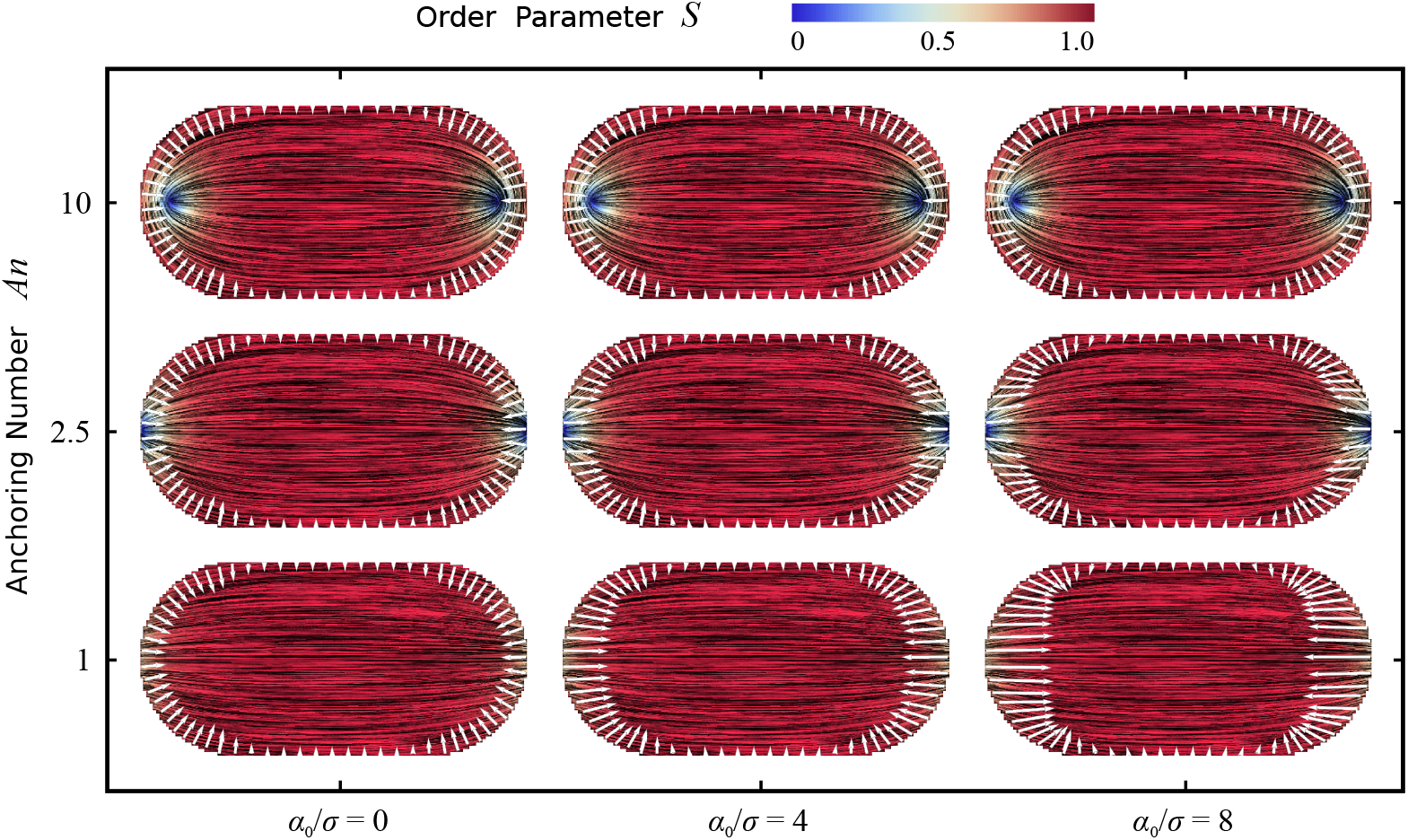
Traction forces of cells adhering to a micropattern in the shape of a stadium (same shape as in Fig 2). The black lines in the interior of the cells represent the orientation field ***n*** = (cos *θ*_SF_, sin *θ*_SF_) of the stress fibers, the background color indicates the nematic order parameter *S*, and the white arrows represents the traction forces exerted on the substrate by the cell. On the vertical axis the anchoring number *An* = *WR/K* = 1, 2.5, 10 (with *R* = 60 lattice sites) is varied, and on the horizontal axis the ratio *α*_0_*/σ* = 0, 4, 8. The spatial averages of the order parameter are given, from left to right, by: 0.92; 0.92; 0.92 (top row), 0.94; 0.94; 0.94 (middle row), and 0.97; 0.97; 0.97 (bottom row). For all cells, *δ* = 9 lattice sites, *λ/σ* = 60 lattice sites, *µ/λ* = 1*/*30 lattice sites, *µ/*|*γ*_ad_| = 1*/*20 of the area of a lattice site, and *K*Δ*t/*(*ξ*_*r*_*R*^2^) = 2.5 *·* 10^*−*6^. For definitions of Δ*t* and *ξ*_*r*_, see Section 4.

The other extreme, where *α*_0_*/σ* = 8 is shown in the right column of Fig 6. For *An* = 10 (top figure) the difference with the case of zero directed contractility is negligible because the stress fibers (***n***) are parallel to the cell boundary (perpendicular to ***N***) everywhere, hence *α*_0_*S*(***n* · *N***)***n*** = **0** [Eq (13)] independent of the value of *α*_0_. For lower anchoring numbers (*An* = 1, *An* = 2.5; middle and bottom figure), however, the effect of increasing *α*_0_ is evident, as the traction forces largely align with the local stress fiber orientation. Because of the longitudinal orientation of the stress fibers, the traction forces at the end of the circular caps increase in magnitude with respect to the forces at other locations, and this effect is strongest for *An* = 1 because in that case stress fibers are almost perpendicular to the ends of the circular caps. Moreover, unlike in many other models [12, 14, 34–36], the traction forces at the caps deviate significantly from the normal vector of the cell boundary. Interestingly, this deviation in orientation from the surface normal is largest for intermediate anchoring number (*An* = 2.5), because the tangential component of the traction force is proportional to (***n* · *N***)(***n* · *T***) [Eq (13)], which is maximized at a *π/*4 angle between the stress fibers and the boundary.

For intermediate values of *α*_0_ (middle column in Fig 6, *α*_0_*/σ* = 4), traction force magnitude and orientation are determined by a combination of local boundary shape and local stress fiber orientation. The resulting traction force configurations for *An* = 1 and *An* = 2.5 qualitatively reproduce experimental traction force patterns of fibroblasts reported by Oakes *et al*. [14] and of endothelial cells reported by Roca-Cusachs *et al*. [15]. Not only are these experimentally observed traction forces larger at the spherical caps than elsewhere, as was already reproduced by the isotropic model put forward in Ref. [14], but they clearly deviate from the normal direction toward the stress fibers, which are oriented along the longitudinal direction of the stadium. Hence, our model qualitatively reproduces not only the structure of the cytoskeleton (see Section 2.2), but also both the magnitudes and the directions of the experimentally observed traction forces.

Next, we study traction forces on an adhesive mircropattern in the shape of a crossbow (see Figs 3b and 5). Fig 7 shows the cells and the traction forces at the cell boundary on the adhesive part of the substrate (Fig 3b) as a function of the anchoring number *An* (vertical axis) and the directed bulk contractility *α*_0_ (horizontal axis). For *α*_0_ = 0 (left column of Fig 7), traction forces are again independent of the stress fiber orientation and normal to the surface. Because of larger local curvature, the forces are larger at the left, bottom, and right sides of the cell. Similar to what we observed in Fig 6, for *An* = 10 the traction forces are independent on the directed contractility *α*_0_ because stress fibers parallel to the cell boundary cannot pull on it. For lower anchoring numbers, increasing the directed bulk contractility is more interesting. First, traction forces are again deviating from the cell boundary’s normal vector, and are biased toward the local stress fiber orientation. This is most evident in the cells with *An* = 1, *An* = 2.5 and *α*_0_*/σ* = 8, where traction forces in the top left and top right of the crossbow point downward and forces in the bottom left and bottom right corners point upward. Moreover, the forces increase in magnitude, as a function of *α*_0_, at the top and bottom of the pattern, where the stress fibers orient perpendicular to the boundary. Similar to what we observed in Fig 6 on the stadiums, the magnitudes of the forces are largest for *An* = 1, whereas the orientations of the forces deviate most from the surface normal at *An* = 2.5.

**Fig 7.**
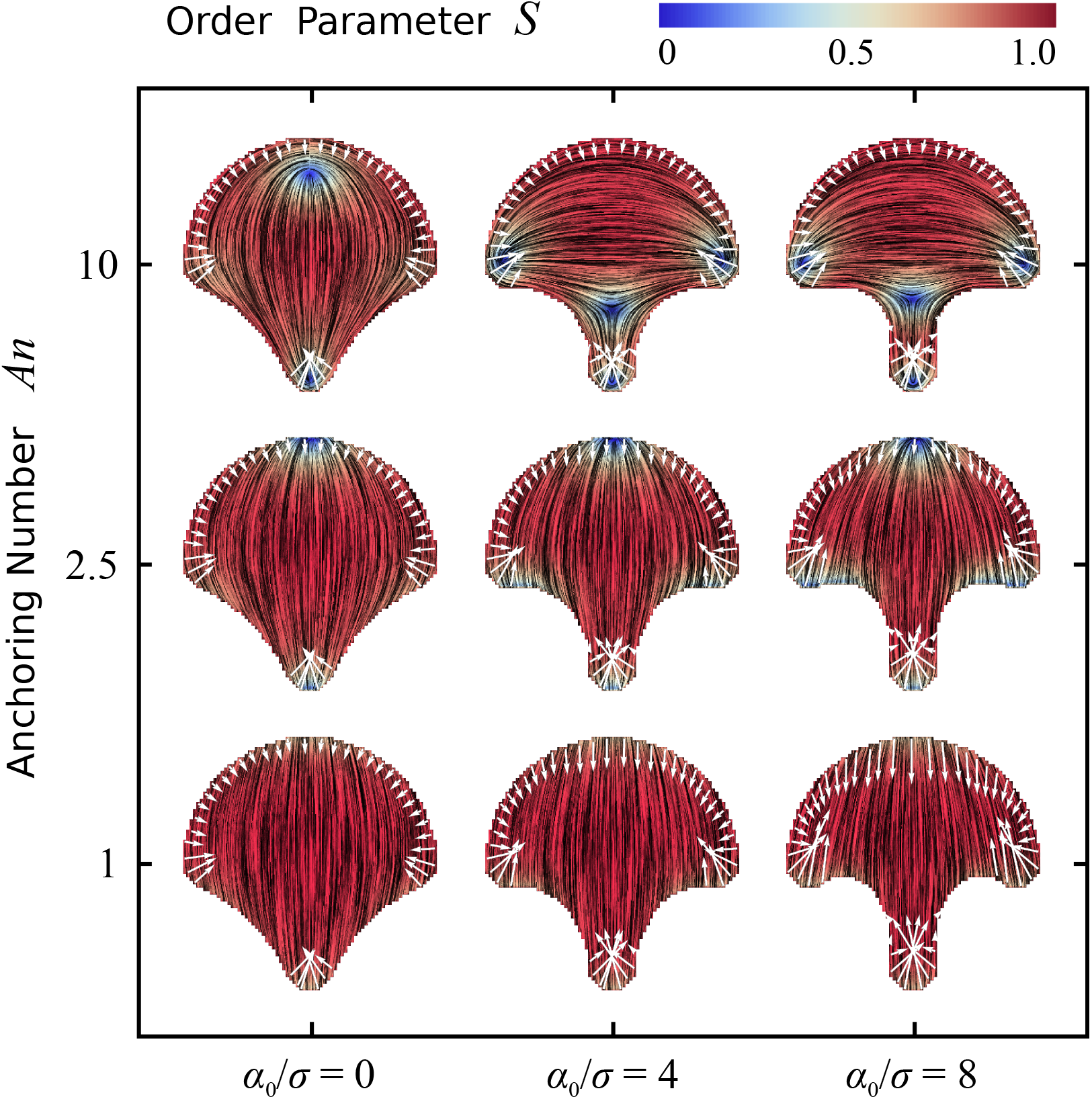
Traction forces of cells adhering to a micropattern in the shape of a crossbow (same shape as in Figs 3b and 5). The black lines in the interior of the cells represent the orientation field ***n*** = (cos *θ*_SF_, sin *θ*_SF_) of the stress fibers, the background color indicates the nematic order parameter *S*, and the white arrows represents the traction forces exerted on the substrate by the cell. On the vertical axis the anchoring number *An* = *WR/K* = 1, 2.5, 10 (with *R* = 60 lattice sites) is varied, and on the horizontal axis the ratio *α*_0_*/σ* = 0, 4, 8. The spatial averages of the order parameter are given, from left to right, by: 0.82; 0.80; 0.81 (top row), 0.88; 0.86; 0.84 (middle row), and 0.94; 0.94; 0.93 (bottom row). For all cells, *δ* = 9 lattice sites, *λ/σ* = 60 lattice sites, *µ/λ* = 1*/*30 lattice sites, *µ/*|*γ*_ad_| = 1*/*20 of the area of a lattice site, and *K*Δ*t/*(*ξ*_*r*_*R*^2^) = 2.5 *·* 10^*−*6^. For definitions of Δ*t* and *ξ*_*r*_, see Section 4.

Comparing the results for low and intermediate anchoring and nonzero directed bulk contractility in Fig 7 with the experimentally reported traction forces of epithelial cells on a crossbow pattern in Tseng *et al*. [6], yields a number of interesting observations. First, the general orientation of the traction forces around the boundary and the increased magnitudes of the traction forces at the bottom, left, and right sides of the pattern agree well between theory and experiment, but these features were previously explained by an isotropic Cellular Potts Model in the work of Albert and Schwarz [12]. However, Tseng *et al*. [6] additionally report an increase in traction force magnitude at the top of the pattern. This increase was not reproduced in the isotropic model of Albert and Schwarz [12], who suggested this discrepancy might be due to the presence of stress fibers along the long side of the pattern (vertical) in the experiments, which areabsent in their model. Our results do show these stress fibers and a resulting increase in traction force magnitude at the top of the pattern, demonstrating that the anisotropy of the actin cytoskeleton is a likely explanation for the discrepancy between the experimental data in Ref. [6] and the simulations in Ref. [12]. Albert and Schwarz further report that forces in the left and right arms of the crossbow pattern are directed more upwards in the experimental data than in their model. The present model does not give better predictions for this observation, possibly due to the fact that the effect of the contraction of internal actin fibers is not well represented in our model.

## 3 Discussion

In this paper we developed a hybrid liquid crystal-Cellular Potts Model (LC-CPM) framework to account for the effect of directed stresses, originating from actin stress fibers, on cell contractility. We first tested the method by comparing the shapes of single cellular arcs in the CPM with exact analytical results. Then, we combined our model for anisotropic contractility with a model for the organization of the actin cytoskeleton [37], where the orientation of stress fibers is governed by a competition between the tendency of stress fibers to align parallel to each other and tangentially with respect to the cell contour. We verified that the hybrid LC-CPM method reproduces earlier numerical results on rectangular cells [37], and qualitatively reproduces experimentally observed stress fiber distributions on convex adhesive micropatterns in the shape of circles [79] and stadiums [14], and on non-convex patterns in the shape of a crossbow [6]. Finally, we calculated the traction forces that cells exert on adhesive micropatterns, and showed that their direction is strongly influenced by the anisotropy of the stress fibers. In particular, we presented a numerical method that, unlike many theoretical models [12, 14, 34, 35], produces traction forces whose orientations deviate away from the normal vector of the cell edge and toward the stress fiber orientation, in agreement with experimental observations [13, 15, 16]. Importantly, our model qualitatively predicts prominent anisotropic features in traction force configurations reported in previously published experimental data on fibroblasts and endothelial cells adhering to micropatterns with stadium shape [14, 15] and epithelial cells on micropatterns with crossbow shape [6], which were not captured by earlier models [12, 14]. Hence, our numerical approach rationalizes findings on anisotropic traction force patterns in earlier experiments, suggesting an important role for stress fibers that is worth investigating, both experimentally and theoretically, in much greater detail.

The main drawback of our current model is that it is *a priori* unclear how to adapt the phenomenological parameters of the cytoskeleton, captured in the anchoring number *An* = *WR/K* [Eq (11)], for different cells types, environmental conditions, or even different arcs within the same cell (see also the discussion of Fig 4 in Section 2.3). For instance, in Ref. [37] we employed this model of the cytoskeleton to study concavely shaped epithelioid and fibroblastoid cells adhering to microfabricated pillar arrays, and found that cells are best described by sufficiently large boundary anchoring of the stress fibers (*An* ≳ 3). In the current study, however, we found qualitative agreement with experimental traction force patterns of fibroblasts, endothelial cells and epithelial cells, adhering to convexly shaped micropatterns, for lower boundary anchoring values (*An* = 1 and *An* = 2.5). A possible explanation for this discrepancy can be found in the fact that concave cell edges promote stress fiber formation more than convex cell edges do [6, 69, 71, 80], an effect that is not present in our current model. We emphasize, however, that our methods for implementing directed cell contractility [Eq (6)] and non-normal traction forces [Eq (13)] in the CPM do not crucially depend on our model for the cytoskeleton (Section 2.2), but can be combined with any model for the orientation of stress fibers. This offers opportunities to, in the future, include other effects into the model, such as the viscoelasticity of stress fibers and actin filament turnover [82, 83], the distinction between different subtypes of stress fibers [84], spatial variations in actin densities [14, 69–71], and the increase of cytoskeletal tension with substrate stiffness [85] or with substrate area [11, 14, 20].

We have focused our traction force analysis on the edge of the cell. Although this is not unreasonable, as traction forces are largest far away from the cell centroid [23, 24, 27] and accumulate at the cell periphery [25, 26], predicting traction forces in the cell interior would help in quantitative comparisons between theory and experiment in the future [12, 86]. Furthermore, our model could serve as a starting point for studying the role of stress fiber contractility and anisotropy in cell spreading and migration. In the framework of the Cellular Potts Model, this could be obtained by combining our model for stress fiber contractility with current CPM implementations of cell motility that depend on the formation of a lamellipodium at the cell’s leading edge [60–62]. Integrating our stress fiber description with these models of cell motility would simulate the persistent motility of cells more realistically, as it would introduce better descriptions of pulling forces at the cell’s rear end [87, 88]. The resulting model could then be employed to computationally study the role of cytoskeletal anisotropy in spreading of single [12, 28, 69, 89] or multiple [90] cells on micropatterns, to explore the effects of cytoskeletal anisotropy on cell-substrate interactions [86, 91], and to better understand the migration of persistent cells in complex topographies [92, 93]. Finally, as the Cellular Potts Model is an excellent tool to computationally study multicellular systems [56], our work can serve as a starting point to study the role of cytoskeletal anisotropy in tissues [66, 94–100].

## 4 Methods

To update the cell shape, we ran the model for a number of timesteps chosen such that the cell shape and the Q-tensor field reached a steady-state configuration. Each timestep consisted of an series of updates of the Q-tensor field, followed by a Monte Carlo step of the cellular Potts model. The simulations in Figure 1 were run for 20,000 timesteps. The simulations in Figures 2-7 were run for 30,000 timesteps, whereas the simulations for *An* = 2.5 required only 10,000 timesteps to steady state.

### Initialization

We first defined the lattice sites that were part of the adhesive micropattern (Figures 1a, 2a, and 3) and the lattice sites that were initially part of the cell. Figure 1a shows the initial cell shape in blue and the adhesive micropattern in grey. In all other figures the initial cell shape was identical to the shape of the adhesive micropattern. The cytoskeleton was initialized by defining the values of 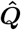 at every lattice site. To this end we selected appropriate values for *S* and *θ*_SF_ and calculated 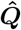 using Eq. (8).

In Figures 1b,c we initialized each lattice site to *S*=0. In Figures 1d-f we set 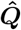 such that *S* = 1 and *θ*_SF_ = *π/*2 at each lattice site. In all the other figures, we assigned random initial values for the orientation − *π/*2 ≤*θ*_SF_ ≤ *π/*2 and the nematic order parameter 0 ≤ *S* ≤ 1. This initial cell configuration does not describe a realistic cell, but it minimizes the risk of a possible bias in the final configuration.

### Cytoskeleton updates

To update the Q-tensor field, the free energy *F*_cyto_ in Eq. (9) was minimized using overdamped relaxational dynamics:

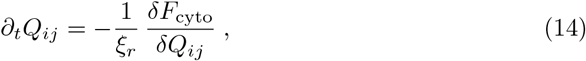

where *Q*_*ij*_ represents the various components of 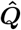, and *ξ*_*r*_ is a rotational friction coefficient required for dimensional consistency. *ξ*_*r*_ dictates the rate of the relaxational dynamics but does not affect the steady-state solution. To ensure the free-energy to be minimal in steady-state (i.e., when *∂*_*t*_*Q*_*ij*_ = 0), Eq. (14) was solved with Neumann boundary conditions:

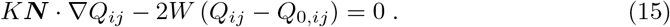

At each time step Δ*t*, 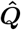 is updated at every lattice point by discretizing Eq. (14) using a forward Euler method. For details on this procedure and on the boundary conditions, see Ref. [37]. For every Monte Carlo step (MCS) of the cellular Potts model, 50 time steps of the cytoskeleton were performed. This step was skipped for the simulations with predefined cytoskeletal configurations shown in Figure 1.

### Cellular Potts model

We used the Cellular Potts model to describe a cell on a micropatterned substrate. The cell is modeled as a collection of lattice sites 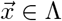 on a regular square lattice Λ*⊂* ℤ^2^, where the lattice sites 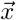 identify the centers of the ‘pixels’ and the arrow notation 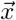 is used specifically to identify discrete lattice coordinates. The lattices are assigned a spin or cell identifier 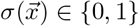, where 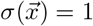 indicates that the lattice site is covered by the cell and 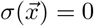 indicates a part of the substrate not covered by the cell. The system evolves through random attempts to retract or extend the cell boundaries, with probabilities depending on the passive and active energy changes in the system (Eq. (7)),

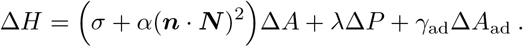

To simulate a random cell extension or retraction, the algorithm iteratively picks a random lattice site 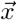, and calculates the energy change Δ*H* resulting from a copy of 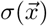 into an adjacent lattice site 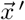. It then accepts this copy with probability depending on the change in energy, Δ*H*,

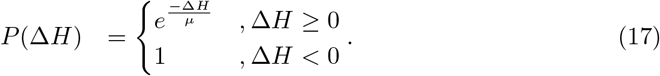

*µ* is a motility parameter, aka ‘cellular temperature’, and represents the amount and magnitude of active, random cell fluctuations, which may act against the energy gradients.

To calculate the length of the cell perimeter, *P*, we employ a previously proposed algorithm that smoothens out lattice effects [101]. Briefly, for every lattice site inside the cell a circular neighborhood of radius *R*, centered at this lattice site, is defined. For all simulations we have used 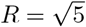. Then, the algorithm counts, for every lattice site, the number of lattice sites whose center lies inside the neighborhood but outside the cell, and the resulting numbers are added up for all lattice sites inside the cell. Finally, to obtain *P* the result is divided by a scaling factor Ξ that depends on the neighborhood size used. In all simulations we have used the numerically determined scaling factor of Ξ = 11 (see Table 2 in Ref. [101]).

The normal vectors at a lattice site 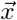 at the cell edge, 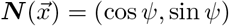, are calculated by again considering a circular neighborhood of radius *R* centered at 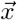, as illustrated in Figure 8. Here the black dot indicates 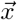, the lattice sites inside the cell are blue, and those outside the cell are white. Similar to Ref. [12], the normal vector 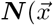 at a given edge lattice site is then calculated as the unit vector pointing from that lattice site to the center of mass (red dot) of the collection of lattice sites that are both inside the cell and inside the neighborhood (cyan lattice sites). In our simulations we have used 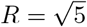.

**Fig 8.**
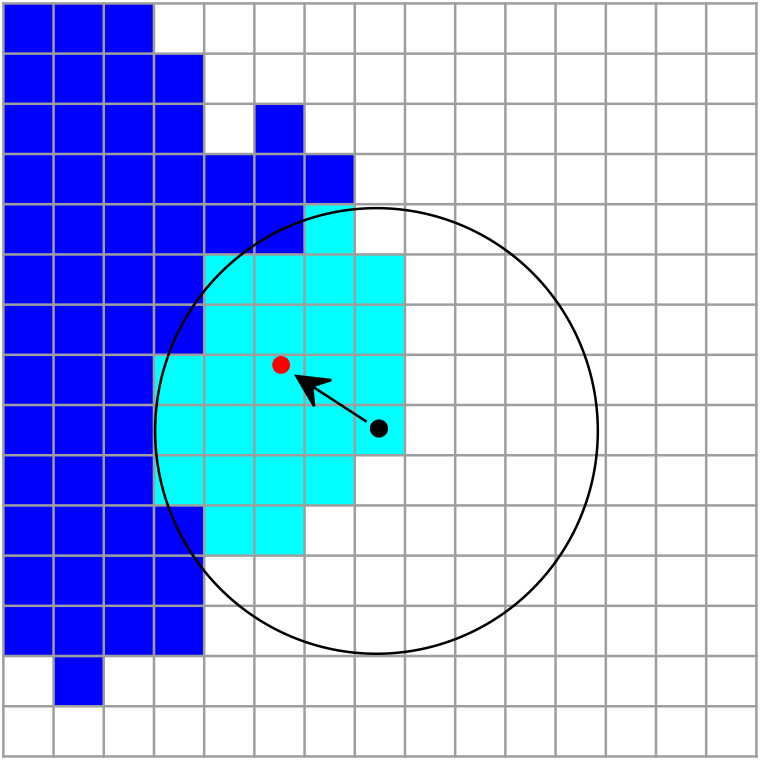
Schematic of the calculation of the normal vector at a lattice site (black dot) at the edge of the cell. The cell is represented by the blue lattice sites. The normal vector is calculated as the unit vector pointing from that lattice site to the center of mass (red dot) of the collection of lattice sites that are both inside the cell and inside a circular neighborhood of radius *R* around that lattice site (cyan lattice sites).

### Final configurations, visualization and calculation of traction forces

The final configurations were determined by averaging the configurations of the last 3,000 MCS of the simulation. If a lattice site was inside the cell for more than 50% of this time, it was considered inside the final configuration. Otherwise, it was considered to be outside the cell. For lattice sites inside the final configuration, the local 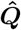 is determined by averaging 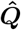 over all MCS within the last 3,000 MCS of the simulation when this lattice site was inside the cell. The cytoskeleton was visualized using Mathematica Version 11.3 (Wolfram Research, Champaign, IL) using the line integral convolution tool. For the visualization step we defined *S* = 1 outside the cell, whereas *θ*_SF_ was undefined outside the cell.

Traction forces were calculated using Eq. (13) and plotted at an interval of five boundary lattice sites. The curvature, *κ*, in this equation is obtained by calculating the derivative of the normal angle *ψ* to the arc length *s, ∂*_*s*_*ψ*. We first define the boundary lattice sites as the set of sites that are inside the cell, and have at least one 8-connected neighboring lattice site outside the cell. We then assign to all boundary lattice sites an ordinal label *i* such that the two 4-connected neighboring boundary lattice sites have labels *i −* 1 and *i* + 1. At each of these boundary lattice sites we calculate *ψ* by calculating the normal vector using a neighborhood with *R* = 10.5. The derivative *∂*_*s*_*ψ* is then estimated using a finite difference approximation,

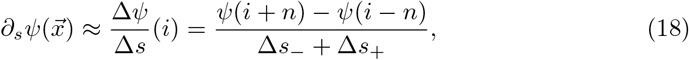

where *i* is the ordinal index of 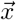, and Δ*s*_*−*_ is the Euclidian distance between lattice sites *i − n* and *i* and Δ*s*_+_ is the Euclidean distance between lattice sites *i* and *i* + *n*. The value of *n* that produces the most accurate approximation of the curvature depends on the actual curvature of the cell boundary. Therefore, we first estimate the radius of curvature *r* = 1*/κ* at a given lattice site using *n* = 3. Then, we calculate the curvature again using *n* = 2 if the initial radius of curvature *r <* 14 lattice sites, *n* = 3 for 14 *< r <* 25 lattice sites, *n* = 4 for 25 *< r <* 35 lattice sites, and *n* = 5 if *r >* 35 lattice sites. In this way we obtain the curvature at each boundary lattice site with a systematic error *<* 1% and a random error *<* 5%.

## Acknowledgments

This work was supported by funds from the Netherlands Organisation for Scientific Research (NWO/OCW), as part of the Frontiers of Nanoscience program (L. G.), the Netherlands Organization for Scientific Research (NWO-ENW) within the Innovational Research Incentives Scheme (L.G., Vidi 2015; R. M. H. M., Vici 2017, No. 865.17.004), and the Leiden/Huygens fellowship (K. S.).

## References

1. Engler AJ, Sen S, Sweeney HL, Discher DE. Matrix elasticity directs stem cell lineage specification. Cell. 2006;126(4):677–689. doi:10.1016/j.cell.2006.06.044.

2. Trappmann B, Gautrot JE, Connelly JT, Strange DGT, Li Y, Oyen ML, et al. Extracellular-matrix tethering regulates stem-cell fate. Nature materials. 2012;11(7):642–649. doi:10.1038/nmat3339.

3. Lo CM, Wang HB, Dembo M, Wang YL. Cell movement is guided by the rigidity of the substrate. Biophysical journal. 2000;79(1):144–152. doi:10.1016/S0006-3495(00)76279-5.

4. Sochol RD, Higa AT, Janairo RRR, Li S, Lin L. Unidirectional mechanical cellular stimuli via micropost array gradients. Soft Matter. 2011;7(10):4606. doi:10.1039/c1sm05163f.

5. Sawada Y, Tamada M, Dubin-Thaler BJ, Cherniavskaya O, Sakai R, Tanaka S, et al. Force sensing by mechanical extension of the Src family kinase substrate p130Cas. Cell. 2006;127(5):1015–26. doi:10.1016/j.cell.2006.09.044.

6. Tseng Q, Wang I, Duchemin-Pelletier E, Azioune A, Carpi N, Gao J, et al. A new micropatterning method of soft substrates reveals that different tumorigenic signals can promote or reduce cell contraction levels. Lab Chip. 2011;11(13):2231–2240. doi:10.1039/C0LC00641F.

7. Kraning-Rush CM, Califano JP, Reinhart-King CA. Cellular Traction Stresses Increase with Increasing Metastatic Potential. PLOS ONE. 2012;7(2):1–10. doi:10.1371/journal.pone.0032572.

8. Peschetola V, Laurent VM, Duperray A, Michel R, Ambrosi D, Preziosi L, et al. Time-dependent traction force microscopy for cancer cells as a measure of invasiveness. Cytoskeleton. 2013;70(4):201–214. doi:10.1002/cm.21100.

9. Pushkarsky I, Tseng P, Black D, France B, Warfe L, Koziol-White CJ, et al. Elastomeric sensor surfaces for high-throughput single-cell force cytometry. Nature Biomedical Engineering. 2018;2(2):124–137. doi:10.1038/s41551-018-0193-2.

10. Bischofs IB, Schmidt SS, Schwarz US. Effect of Adhesion Geometry and Rigidity on Cellular Force Distributions. Physical Review Letters. 2009;103(4):1–4. doi:10.1103/PhysRevLett.103.048101.

11. Rape AD, Guo WH, Wang YL. The regulation of traction force in relation to cell shape and focal adhesions. Biomaterials. 2011;32(8):2043–2051. doi:https://doi.org/10.1016/j.biomaterials.2010.11.044.

12. Albert PJ, Schwarz US. Dynamics of Cell Shape and Forces on Micropatterned Substrates Predicted by a Cellular Potts Model. Biophysical Journal. 2014;106(11):2340–2352. doi:https://doi.org/10.1016/j.bpj.2014.04.036.

13. Mandal K, Wang I, Vitiello E, Orellana LAC, Balland M. Cell dipole behaviour revealed by ECM sub-cellular geometry. Nature Communications. 2014;5:5749.

14. Oakes PW, Banerjee S, Marchetti MC, Gardel ML. Geometry Regulates Traction Stresses in Adherent Cells. Biophysical Journal. 2014;107(4):825–833. doi:https://doi.org/10.1016/j.bpj.2014.06.045.

15. Roca-Cusachs P, Alcaraz J, Sunyer R, Samitier J, Farré R, Navajas D. Micropatterning of Single Endothelial Cell Shape Reveals a Tight Coupling between Nuclear Volume in G1 and Proliferation. Biophysical Journal. 2008;94(12):4984–4995. doi:https://doi.org/10.1529/biophysj.107.116863.

16. Pomp W, Schakenraad K, Balcioğlu HE, van Hoorn H, Danen EHJ, Merks RMH, et al. Cytoskeletal Anisotropy Controls Geometry and Forces of Adherent Cells. Physical Review Letters. 2018;121(17):178101. doi:10.1103/PhysRevLett.121.178101.

17. Tan JL, Tien J, Pirone DM, Gray DS, Bhadriraju K, Chen CS. Cells lying on a bed of microneedles: an approach to isolate mechanical force. Proceedings of the National Academy of Sciences of the United States of America. 2003;100(4):1484–9. doi:10.1073/pnas.0235407100.

18. Reinhart-King CA, Dembo M, Hammer DA. The Dynamics and Mechanics of Endothelial Cell Spreading. Biophysical Journal. 2005;89(1):676–689. doi:https://doi.org/10.1529/biophysj.104.054320.

19. Tolić-Nørrelykke IM, Wang N. Traction in smooth muscle cells varies with cell spreading. Journal of Biomechanics. 2005;38(7):1405–1412. doi:https://doi.org/10.1016/j.jbiomech.2004.06.027.

20. Hanke J, Probst D, Zemel A, Schwarz US, Köster S. Dynamics of force generation by spreading platelets. Soft Matter. 2018;14(31):6571–6581. doi:10.1039/C8SM00895G.

21. Wang N, Ostuni E, Whitesides GM, Ingber DE. Micropatterning tractional forces in living cells. Cell Motility. 2002;52(2):97–106. doi:10.1002/cm.10037.

22. Califano JP, Reinhart-King CA. Substrate Stiffness and Cell Area Predict Cellular Traction Stresses in Single Cells and Cells in Contact. Cellular and Molecular Bioengineering. 2010;3(1):68–75. doi:10.1007/s12195-010-0102-6.

23. Roux C, Duperray A, Laurent VM, Michel R, Peschetola V, Verdier C, et al. Prediction of traction forces of motile cells. Interface Focus. 2016;6(5):20160042. doi:10.1098/rsfs.2016.0042.

24. Lemmon CA, Romer LH. A Predictive Model of Cell Traction Forces Based on Cell Geometry. Biophysical Journal. 2010;99(9):L78–L80. doi:https://doi.org/10.1016/j.bpj.2010.09.024.

25. Riveline D, Zamir E, Balaban NQ, Schwarz US, Ishizaki T, Narumiya S, et al. Focal contacts as mechanosensors: externally applied local mechanical force induces growth of focal contacts by an mDia1-dependent and ROCK-independent mechanism. The Journal of cell biology. 2001;153(6):1175–1186.

26. Balaban NQ, Schwarz US, Riveline D, Goichberg P, Tzur G, Sabanay I, et al. Force and focal adhesion assembly: a close relationship studied using elastic micropatterned substrates. Nature Cell Biology. 2001;3(5):466–472. doi:10.1038/35074532.

27. Edwards CM, Schwarz US. Force Localization in Contracting Cell Layers. Physical Review Letters. 2011;107(12):128101. doi:10.1103/PhysRevLett.107.128101.

28. Kim MC, Neal DM, Kamm RD, Asada HH. Dynamic Modeling of Cell Migration and Spreading Behaviors on Fibronectin Coated Planar Substrates and Micropatterned Geometries. PLOS Computational Biology. 2013;9(2):1–13. doi:10.1371/journal.pcbi.1002926.

29. Bangasser BL, Odde DJ. Master Equation-Based Analysis of a Motor-Clutch Model for Cell Traction Force. Cellular and Molecular Bioengineering. 2013;6(4):449–459. doi:10.1007/s12195-013-0296-5.

30. Lin S, Lampi MC, Reinhart-King CA, Tsui G, Wang J, Nelson CA, et al. Eigenstrain as a mechanical set-point of cells. Biomechanics and Modeling in Mechanobiology. 2018;17(4):951–959. doi:10.1007/s10237-018-1004-0.

31. Oakes PW. Balancing forces in migration. Current Opinion in Cell Biology. 2018;54:43–49. doi:https://doi.org/10.1016/j.ceb.2018.04.006.

32. Banerjee S, Marchetti MC. Controlling cell–matrix traction forces by extracellular geometry. New Journal of Physics. 2013;15(3):035015. doi:10.1088/1367-2630/15/3/035015.

33. Banerjee S, Sknepnek R, Marchetti MC. Optimal shapes and stresses of adherent cells on patterned substrates. Soft Matter. 2014;10:2424–2430. doi:10.1039/C3SM52647J.

34. Ben-Yaakov D, Golkov R, Shokef Y, Safran SA. Response of adherent cells to mechanical perturbations of the surrounding matrix. Soft Matter. 2015;11(7):1412–1424. doi:10.1039/C4SM01817F.

35. Golkov R, Shokef Y. Elastic interactions between anisotropically contracting circular cells. Physical Review E. 2019;99(3):032418. doi:10.1103/PhysRevE.99.032418.

36. Rens EG, Edelstein-Keshet L. From energy to cellular forces in the Cellular Potts Model: An algorithmic approach. PLOS Computational Biology. 2019;15(12):1–23. doi:10.1371/journal.pcbi.1007459.

37. Schakenraad K, Ernst J, Pomp W, Danen EHJ, Merks RMH, Schmidt T, et al. Mechanical interplay between cell shape and actin cytoskeleton organization. Soft Matter. 2020;16(27):6328–6343. doi:10.1039/D0SM00492H.

38. Théry M. Micropatterning as a tool to decipher cell morphogenesis and functions. Journal of Cell Science. 2010;123(24):4201–4213. doi:10.1242/jcs.075150.

39. Schwarz US, Safran SA. Physics of adherent cells. Reviews of Modern Physics. 2013;85(3):1327–1381. doi:10.1103/RevModPhys.85.1327.

40. Trichet L, Le Digabel J, Hawkins RJ, Vedula SRK, Gupta M, Ribrault C, et al. Evidence of a large-scale mechanosensing mechanism for cellular adaptation to substrate stiffness. Proceedings of the National Academy of Sciences. 2012;109(18):6933–6938. doi:10.1073/pnas.1117810109.

41. Van Hoorn H, Harkes R, Spiesz EM, Storm C, Van Noort D, Ladoux B, et al. The nanoscale architecture of force-bearing focal adhesions. Nano Letters. 2014;14(8):4257–4262. doi:10.1021/nl5008773.

42. Dembo M, Wang YL. Stresses at the Cell-to-Substrate Interface during Locomotion of Fibroblasts. Biophysical Journal. 1999;76(4):2307–2316. doi:https://doi.org/10.1016/S0006-3495(99)77386-8.

43. Butler JP, Tolić-Nørrelykke IM, Fabry B, Fredberg JJ. Traction fields, moments, and strain energy that cells exert on their surroundings. American Journal of Physiology-Cell Physiology. 2002;282(3):C595–C605. doi:10.1152/ajpcell.00270.2001.

44. Sabass B, Gardel ML, Waterman CM, Schwarz US. High Resolution Traction Force Microscopy Based on Experimental and Computational Advances. Biophysical Journal. 2008;94(1):207–220. doi:https://doi.org/10.1529/biophysj.107.113670.

45. Burridge K, Chrzanowska-Wodnicka M. Focal Adhesions, Contractility, and Signaling. Annual Review of Cell and Developmental Biology. 1996;12(1):463–519. doi:10.1146/annurev.cellbio.12.1.463.

46. Bar-Ziv R, Tlusty T, Moses E, Safran Sa, Bershadsky a. Pearling in cells: a clue to understanding cell shape. Proceedings of the National Academy of Sciences of the United States of America. 1999;96(18):10140–5. doi:10.1073/pnas.96.18.10140.

47. Bischofs IB, Klein F, Lehnert D, Bastmeyer M, Schwarz US. Filamentous Network Mechanics and Active Contractility Determine Cell and Tissue Shape. Biophysical Journal. 2008;95(7):3488–3496. doi:10.1529/biophysj.108.134296.

48. Banerjee S, Giomi L. Polymorphism and bistability in adherent cells. Soft Matter. 2013;9(21):5251–5260. doi:10.1039/C3SM27791G.

49. Giomi L. Softly constrained films. Soft Matter. 2013;9(34):8121. doi:10.1039/c3sm50484k.

50. Giomi L. Contour models of cellular adhesion. In: Zapperi S, La Porta CAM, editors. Cell migrations: causes and function. Berlin: Springer; 2019. p. in press.

51. Pellegrin S, Mellor H. Actin stress fibres. Journal of cell science. 2007;120(20):3491–3499. doi:10.1242/jcs.018473.

52. Burridge K, Wittchen ES. The tension mounts: Stress fibers as force-generating mechanotransducers. The Journal of Cell Biology. 2013;200(1):9–19. doi:10.1083/jcb.201210090.

53. Pedley TJ, Kessler JO. Hydrodynamic Phenomena in Suspensions of Swimming Microorganisms. Annual Review of Fluid Mechanics. 1992;24(1):313–358. doi:10.1146/annurev.fl.24.010192.001525.

54. Simha RA, Ramaswamy S. Hydrodynamic Fluctuations and Instabilities in Ordered Suspensions of Self-Propelled Particles. Physical Review Letters. 2002;89(5):058101. doi:10.1103/PhysRevLett.89.058101.

55. Glazier JA, Graner F. Simulation of the differential adhesion driven rearrangement of biological cells. Phys Rev E. 1993;47:2128–2154. doi:10.1103/PhysRevE.47.2128.

56. Anderson A, Rejniak K. Single-cell-based models in biology and medicine. Springer Science & Business Media; 2007.

57. Hirashima T, Rens EG, Merks RMH. Cellular Potts modeling of complex multicellular behaviors in tissue morphogenesis. Development, Growth & Differentiation. 2017;59(5):329–339. doi:10.1111/dgd.12358.

58. Vianay B, Käfer J, Planus E, Block M, Graner F, Guillou H. Single Cells Spreading on a Protein Lattice Adopt an Energy Minimizing Shape. Phys Rev Lett. 2010;105(12):128101. doi:10.1103/PhysRevLett.105.128101.

59. Albert PJ, Schwarz US. Modeling cell shape and dynamics on micropatterns. Cell Adhesion & Migration. 2016;10(5):516–528. doi:10.1080/19336918.2016.1148864.

60. Marée AFM, Grieneisen VA, Edelstein-Keshet L. How Cells Integrate Complex Stimuli: The Effect of Feedback from Phosphoinositides and Cell Shape on Cell Polarization and Motility. PLOS Computational Biology. 2012;8(3):1–20. doi:10.1371/journal.pcbi.1002402.

61. Niculescu I, Textor J, de Boer RJ. Crawling and Gliding: A Computational Model for Shape-Driven Cell Migration. PLOS Computational Biology. 2015;11(10):1–22. doi:10.1371/journal.pcbi.1004280.

62. van Steijn L, Wortel IMN, Sire C, Dupré L, Theraulaz G, Merks RMH. Computational modelling of cell motility modes emerging from cell-matrix adhesion dynamics. PLOS Computational Biology. 2022;18(2):1–28. doi:10.1371/journal.pcbi.1009156.

63. Newman M, Barkema G. Monte carlo methods in statistical physics. Oxford University Press: New York, USA; 1999.

64. Ramaswamy S. Active matter. Journal of Statistical Mechanics: Theory and Experiment. 2017;2017(5):054002. doi:10.1088/1742-5468/aa6bc5.

65. Savill NJ, Hogeweg P. Modelling Morphogenesis: From Single Cells to Crawling Slugs. Journal of Theoretical Biology. 1997;184(3):229–235. doi:10.1006/jtbi.1996.0237.

66. van Oers RFM, Rens EG, LaValley DJ, Reinhart-King CA, Merks RMH. Mechanical Cell-Matrix Feedback Explains Pairwise and Collective Endothelial Cell Behavior In Vitro. PLOS Computational Biology. 2014;10(8):1–14. doi:10.1371/journal.pcbi.1003774.

67. de Gennes PG, Prost J. The Physics of Liquid Crystals. International Series of Monogr. Clarendon Press; 1995. Available from: https://books.google.nl/books?id=0Nw-dzWz5agC.

68. D Pathak A, Deshpande VS, McMeeking RM, Evans AG. The simulation of stress fibre and focal adhesion development in cells on patterned substrates. Journal of The Royal Society Interface. 2008;5(22):507–524. doi:10.1098/rsif.2007.1182.

69. Théry M, Pépin A, Dressaire E, Chen Y, Bornens M. Cell distribution of stress fibres in response to the geometry of the adhesive environment. Cell Motility. 2006;63(6):341–355. doi:10.1002/cm.20126.

70. Théry M, Racine V, Piel M, Pépin A, Dimitrov A, Chen Y, et al. Anisotropy of cell adhesive microenvironment governs cell internal organization and orientation of polarity. Proceedings of the National Academy of Sciences. 2006;103(52):19771–19776. doi:10.1073/pnas.0609267103.

71. James J, Goluch ED, Hu H, Liu C, Mrksich M. Subcellular curvature at the perimeter of micropatterned cells influences lamellipodial distribution and cell polarity. Cell Motility. 2008;65(11):841–852. doi:10.1002/cm.20305.

72. Vignaud T, Blanchoin L, Théry M. Directed cytoskeleton self-organization. Trends in Cell Biology. 2012;22(12):671–682. doi:https://doi.org/10.1016/j.tcb.2012.08.012.

73. Ladoux B, Mège RM, Trepat X. Front–Rear Polarization by Mechanical Cues: From Single Cells to Tissues. Trends in Cell Biology. 2016;26(6):420–433. doi:https://doi.org/10.1016/j.tcb.2016.02.002.

74. Lam NT, Muldoon TJ, Quinn KP, Rajaram N, Balachandran K. Valve interstitial cell contractile strength and metabolic state are dependent on its shape. Integrative Biology. 2016;8(10):1079–1089. doi:10.1039/c6ib00120c.

75. Gupta SK, Li Y, Guo M. Anisotropic mechanics and dynamics of a living mammalian cytoplasm. Soft Matter. 2019;15(2):190–199. doi:10.1039/C8SM01708E.

76. Nobili M, Durand G. Disorientation-induced disordering at a nematic-liquid-crystal–solid interface. Phys Rev A. 1992;46(10):R6174–R6177. doi:10.1103/PhysRevA.46.R6174.

77. Kleman M, Lavrentovich OD. Topological point defects in nematic liquid crystals. Philosophical Magazine. 2006;86(25-26):4117–4137.

78. Tee YH, Shemesh T, Thiagarajan V, Hariadi RF, Anderson KL, Page C, et al. Cellular chirality arising from the self-organization of the actin cytoskeleton. Nature Cell Biology. 2015;17:445.

79. Jalal S, Shi S, Acharya V, Huang RYJ, Viasnoff V, Bershadsky AD, et al. Actin cytoskeleton self-organization in single epithelial cells and fibroblasts under isotropic confinement. Journal of Cell Science. 2019;132(5). doi:10.1242/jcs.220780.

80. Kilian KA, Bugarija B, Lahn BT, Mrksich M. Geometric cues for directing the differentiation of mesenchymal stem cells. Proceedings of the National Academy of Sciences. 2010;107(11):4872–4877. doi:10.1073/pnas.0903269107.

81. Labouesse C, Verkhovsky AB, Meister JJ, Gabella C, Vianay B. Cell Shape Dynamics Reveal Balance of Elasticity and Contractility in Peripheral Arcs. Biophysical Journal. 2015;108(10):2437–2447. doi:10.1016/j.bpj.2015.04.005.

82. D Étienne J, Fouchard J, Mitrossilis D, Bufi N, Durand-Smet P, Asnacios A. Cells as liquid motors: Mechanosensitivity emerges from collective dynamics of actomyosin cortex. Proceedings of the National Academy of Sciences. 2015;112(9):2740–2745. doi:10.1073/pnas.1417113112.

83. Oakes PW, Wagner E, Brand CA, Probst D, Linke M, Schwarz US, et al. Optogenetic control of RhoA reveals zyxin-mediated elasticity of stress fibres. Nature Communications. 2017;8:15817.

84. Lee S, Kassianidou E, Kumar S. Actomyosin stress fiber subtypes have unique viscoelastic properties and roles in tension generation. Molecular Biology of the Cell. 2018;29(16):1992–2004. doi:10.1091/mbc.E18-02-0106.

85. Fu J, Wang YK, Yang MT, Desai RA, Yu X, Liu Z, et al. Mechanical regulation of cell function with geometrically modulated elastomeric substrates. Nature methods. 2010;7(9):733–736. doi:10.1038/nmeth.1487.

86. EG, Merks RMH. Cell Shape and Durotaxis Explained from Cell-Extracellular Matrix Forces and Focal Adhesion Dynamics. iScience. 2020;23(9):101488. doi:10.1016/j.isci.2020.101488.

87. Barnhart E, Lee KC, Allen GM, Theriot JA, Mogilner A. Balance between cell-substrate adhesion and myosin contraction determines the frequency of motility initiation in fish keratocytes. Proceedings of the National Academy of Sciences. 2015;112(16):5045–5050. doi:10.1073/pnas.1417257112.

88. Abaurrea-Velasco C, Auth T, Gompper G. Self-organized motility of vesicles with internal active filaments: self-organized propulsion controls shape, motility, and dynamical response. New Journal of Physics. 2019;21:123024. doi:10.1088/1367-2630/ab5c70.

89. Loosli Y, Luginbuehl R, Snedeker JG. Cytoskeleton reorganization of spreading cells on micro-patterned islands: a functional model. Philosophical Transactions of the Royal Society A: Mathematical, Physical and Engineering Sciences. 2010;368(1920):2629–2652. doi:10.1098/rsta.2010.0069.

90. Albert PJ, Schwarz US. Dynamics of Cell Ensembles on Adhesive Micropatterns: Bridging the Gap between Single Cell Spreading and Collective Cell Migration. PLOS Computational Biology. 2016;12(4):1–34. doi:10.1371/journal.pcbi.1004863.

91. Deshpande VS, Mrksich M, McMeeking RM, Evans AG. A bio-mechanical model for coupling cell contractility with focal adhesion formation. Journal of the Mechanics and Physics of Solids. 2008;56(4):1484–1510. doi:https://doi.org/10.1016/j.jmps.2007.08.006.

92. Wondergem JAJ, Mytiliniou M, de Wit FCH, Reuvers TGA, Holcman D, Heinrich D. Chemotaxis and topotaxis add vectorially for amoeboid cell migration. bioRxiv. 2019;doi:10.1101/735779.

93. Schakenraad K, Ravazzano L, Sarkar N, Wondergem JAJ, Merks RMH, Giomi L. Topotaxis of active Brownian particles. Phys Rev E. 2020;101:032602. doi:10.1103/PhysRevE.101.032602.

94. Eastwood M, Mudera VC, McGrouther DA, Brown RA. Effect of precise mechanical loading on fibroblast populated collagen lattices: morphological changes. Cell Motility and the Cytoskeleton. 1998;40(1):13–21. doi:10.1002/(SICI)1097-0169(1998)40:1¡13::AID-CM2¿3.0.CO;2-G.

95. Van der Schaft DWJ, Van Spreeuwel ACC, Van Assen HC, Baaijens FPT. Mechanoregulation of Vascularization in Aligned Tissue-Engineered Muscle: A Role for Vascular Endothelial Growth Factor. Tissue Engineering Part A. 2011;17(21-22):2857–2865. doi:10.1089/ten.tea.2011.0214.

96. Wartlick O, Mumcu P, Jülicher F, Gonzalez-Gaitan M. Understanding morphogenetic growth control - lessons from flies. Nature Reviews Molecular Cell Biology. 2011;12(9):594–604. doi:10.1038/nrm3169.

97. Santos-Oliveira P, Correia A, Rodrigues T, Ribeiro-Rodrigues TM, Matafome P, Rodríguez-Manzaneque JC, et al. The Force at the Tip - Modelling Tension and Proliferation in Sprouting Angiogenesis. PLoS Computational Biology. 2015;11(8):1–20. doi:10.1371/journal.pcbi.1004436.

98. Vijayraghavan DS, Davidson LA. Mechanics of neurulation: From classical to current perspectives on the physical mechanics that shape, fold, and form the neural tube. Birth Defects Research. 2017;109(2):153–168. doi:10.1002/bdra.23557.

99. Saw TB, Doostmohammadi A, Nier V, Kocgozlu L, Thampi S, Toyama Y, et al. Topological defects in epithelia govern cell death and extrusion. Nature. 2017;544:212.

100. Barton DL, Henkes S, Weijer CJ, Sknepnek R. Active Vertex Model for cell-resolution description of epithelial tissue mechanics. PLOS Computational Biology. 2017;13(6):1–34. doi:10.1371/journal.pcbi.1005569.

101. Magno R, Grieneisen VA, Marée AF. The biophysical nature of cells: potential cell behaviours revealed by analytical and computational studies of cell surface mechanics. BMC biophysics. 2015;8(1):8.

